# A non-canonical microRNA derived from the snaR-A non-coding RNA targets a metastasis inhibitor

**DOI:** 10.1101/2021.03.20.436234

**Authors:** Daniel Stribling, Yi Lei, Casey M. Guardia, Lu Li, Christopher J. Fields, Pawel Nowialis, Rene Opavsky, Rolf Renne, Mingyi Xie

## Abstract

MicroRNAs (miRNAs) are small noncoding RNAs that function as critical post-transcriptional regulators in various biological processes. While most miRNAs are generated from processing of long primary transcripts via sequential Drosha and Dicer cleavage, some miRNAs that bypass Drosha cleavage can be transcribed as part of another small non-coding RNA. Here, we develop the <u>T</u>arget-<u>O</u>riented <u>mi</u>RNA <u>D</u>iscovery (TOMiD) bioinformatic analysis method to identify Drosha-independent miRNAs from <u>A</u>r<u>g</u>onaute <u>c</u>rosslinking <u>a</u>nd <u>s</u>equencing of <u>h</u>ybrids (Ago-CLASH) datasets. Using this technique, we discovered a novel miRNA derived from a primate specific non-coding RNA, the small NF90 associated RNA A (snaR-A). The miRNA derived from snaR-A (miR-snaR) arises independent of Drosha processing, but requires Exportin-5 and Dicer for biogenesis. We identify that miR-snaR is concurrently upregulated with the full snaR-A transcript in cancer cells. Functionally, miR-snaR associates with Ago proteins and targets NME1, a key metastasis inhibitor, contributing to snaR-A’s role in promoting cancer cell migration. Our findings suggest a functional link between a novel miRNA and its precursor non-coding RNA.

## INTRODUCTION

MicroRNAs (miRNAs) are ∼22 nucleotide (nt) RNAs that serve an essential regulatory role via targeted silencing and degradation of messenger RNAs (mRNAs) (Bartel 2018). MiRNA targeting is primarily mediated by Watson-Crick base-pairing between the miRNA seed region (nts 2-7) and a complimentary target sequence often located within the mRNA 3’ untranslated region (UTR). The pervasive influence of miRNA-driven regulation is observed in nearly all physiological processes, including cell growth, stress responses, metabolism, and apoptosis, with a predicted ≥70% of human mRNAs regulated by miRNAs (Miranda et al. 2006). Given the major role played by miRNAs in normal cellular function, disruption of miRNA expression can lead to significant developmental and physiological defects and miRNA dysregulation has been shown to be a significant driver of carcinogenesis (Bartel 2018). As such, both uncovering the mechanisms of miRNA biogenesis and identifying miRNA targets are crucial for the understanding of intrinsic cellular regulatory processes and for designing effective therapeutics of diseases associated with miRNA dysregulation.

Since the initial discovery of miRNAs in *C. elegans*, the canonical miRNA biogenesis pathway has been well-characterized (Ha and Kim 2014). Canonical miRNA processing begins with transcription of primary miRNA transcripts (pri-miRNAs), most often by RNA polymerase II (Pol II) (Lee et al. 2002), followed by nuclear cleavage via the Microprocessor complex (composed of Drosha and DGCR8) (Nguyen et al. 2015). The resulting precursor miRNA (pre-miRNA) hairpins are then exported to the cytoplasm by Exportin 5 (XPO5) in a Ran-GTP-dependent manner (Lund et al. 2004). In the cytoplasm, Dicer cleaves pre-miRNA hairpins producing ∼22 nt mature miRNA molecules that tightly associate with a member of the Argonaute (Ago) family of proteins (Bernstein et al. 2001). This Ago-miRNA complex is known as the RNA-induced silencing complex (RISC) and acts to mediate mRNA stability and translation as a part of the post-transcriptional regulatory network (Kawamata and Tomari 2010).

In addition to the canonical pathway, alternative miRNA biogenesis mechanisms bypassing Drosha or Dicer cleavage have been identified (Xie and Steitz 2014). The first identified Drosha-independent miRNAs are mirtrons, which are pre-miRNAs that arise as products of splicing instead of via Drosha cleavage (Ruby et al. 2007; Okamura et al. 2008). Another group of Drosha-independent miRNAs originate from short transcripts derived immediately downstream of Pol II transcription start sites (TSS) (Babiarz et al. 2008; Zamudio et al. 2014). These pre-miRNAs contain a 7-methylguanine cap at the 5’ terminus and are therefore formed independently of Drosha (Xie et al. 2013; Sheng et al. 2018). Additionally, Drosha-independent miRNAs can be processed from other small non-coding RNAs (ncRNAs), including small nucleolar RNAs (snoRNAs) and transfer RNAs (tRNAs) (Babiarz et al. 2008; Ender et al. 2008; Lemus-Diaz et al. 2020). In the absence of Drosha, these non-canonical miRNAs are upregulated in cells, presumably because they are free of competition from canonical miRNAs for access to the limited cellular miRNA machinery (Kim et al. 2016; Sheng et al. 2018). Recent studies have demonstrated the increasingly-emerging functions of these non-canonical miRNAs in several diseases, including cancer (Stavast and Erkeland 2019).

Although miRNA biogenesis pathways have been well characterized, miRNA target identification remains a major challenge. Recent methodological developments in miRNA sequence analysis have significantly improved the accuracy of miRNA target prediction on the genome-wide scale (Lewis et al. 2005; Agarwal et al. 2015). However, these predictions rely heavily on conserved base pairing of the miRNA seed region, which can lead to a high error rate given the high frequency of 6-nt seed-matches in the transcriptome (Pinzon et al. 2017) and the existence of other functional non-seed interactions between miRNAs and their targets (Broughton et al. 2016; Zhang et al. 2018). Biochemical approaches such as high-throughput sequencing of RNAs isolated by crosslinking immunoprecipitation (HITS-CLIP) provide a powerful platform to identify RNA-protein interactions *in vivo* (Licatalosi et al. 2008; Darnell 2010). In this method, <u>u</u>ltra<u>v</u>iolet (UV) irradiation is used to covalently crosslink interacting RNA and protein molecules within cells. As such, Ago HITS-CLIP allows identification of bound miRNAs and their associated mRNAs simultaneously, but relies on assumption-driven analysis methods for determination of miRNA/target pairs (Chi et al. 2009). HITS-CLIP has inspired a number of related methods for direct investigation of miRNA-mRNA targeting, including: <u>C</u>rosslinking, <u>L</u>igation, <u>a</u>nd <u>S</u>equencing of <u>H</u>ybrids (CLASH) (Helwak et al. 2013); <u>Q</u>uick CLASH (qCLASH) (Gay et al. 2018); and <u>C</u>ovalent <u>L</u>igation of <u>E</u>ndogenous <u>A</u>rgonaute-bound <u>R</u>NAs with <u>C</u>rosslinking and <u>I</u>mmuno<u>p</u>recipitation (CLEAR-CLIP) (Moore et al. 2015). These methods induce inter-molecular miRNA-mRNA ligation on the Ago protein, thus producing miRNA-mRNA hybrid molecules for subsequent high-throughput sequencing. As opposed to traditional CLIP methods in which sequenced miRNAs and targets must then be analytically assigned, the hybrid reads generated by CLASH allow unambiguous pairing between miRNAs and their *in vivo* targets. While these methods have been effectively employed for miRNA target identification in several studies, we were interested in determining whether CLASH data could be used for identification of previously unknown miRNA sequences, and specifically for identification of new miRNAs arising via noncanonical biogenesis pathways.

Previous methods developed for analyzing CLASH hybrid sequence data require an annotated database of possible miRNAs and therefore do not permit identification of novel miRNAs sequences. To enable novel miRNA discovery from CLASH data, we developed the <u>T</u>arget-<u>O</u>riented <u>mi</u>RNA <u>D</u>iscovery (TOMiD) analysis method to identify miRNA sequences within reads generated from CLASH experiments by a stepwise process of hybrid read identification independent of previous miRNA annotation. We applied the TOMiD method to search for Drosha-independent miRNAs in Ago-qCLASH datasets obtained from wildtype (WT), Drosha-knockout (KO), and Dicer-KO HCT116 colorectal cancer cell lines (Fields and Xie, manuscript in preparation). We successfully identified a novel Drosha-independent miRNA: miR-snaR, which arises from the small NF90-associated RNA A (snaR-A) family of transcripts (Parrott and Mathews 2007) . We found that miR-snaR biogenesis is Drosha-independent, but dependent on XPO5 and Dicer. In addition, we detected Ago-bound miR-snaR in cells expressing high levels of snaR-A, which is upregulated in multiple cancer-derived cell lines and tumors, including breast, colorectal, liver, and ovarian cancers (Parrott and Mathews 2007; Lee et al. 2016; Lee et al. 2017; Huang et al. 2018; Shi et al. 2019). To investigate the potential function of this miRNA, we identified miR-snaR/target hybrids in multiple Ago-CLASH datasets and discovered that miR-snaR represses expression of non-metastatic cells 1 (NME1), which correspondingly increases mobility of cancer cells.

## RESULTS

### TOMiD identifies Drosha-independent miRNAs in Ago-CLASH datasets

Existing methods for Ago-CLASH data analysis rely on previous sequence annotations to identify miRNA/target hybrids (Travis et al. 2014). To identify novel miRNAs within CLASH data, we developed the TOMiD analysis method (Fig. 1). In this approach, sequence reads generated from a CLASH experiment are first aligned to a reference transcript database. Reads that do not align completely to a single reference transcript are then identified as potential chimeric sequences (hybrids). Potential hybrids are then analyzed via several successive criteria to select for sequences matching characteristics of miRNA/target RNA hybrids specifically. This includes selection based on a specific length of the unaligned portion of the hybrid and on a minimum predicted folding energy of the hybrid via intra-read base-pairing. The unaligned region of each remaining read is then assigned as a candidate miRNA (cmiRNA). In this annotation-naïve approach, a cmiRNA can represent either a previously-known miRNA, a novel miRNA, or a non-miRNA sequence. To maximize the ratio of identified miRNA sequences to nonspecific sequences, several processing steps are then utilized to increase detection of miRNA-containing hybrids.

**Figure 1.**
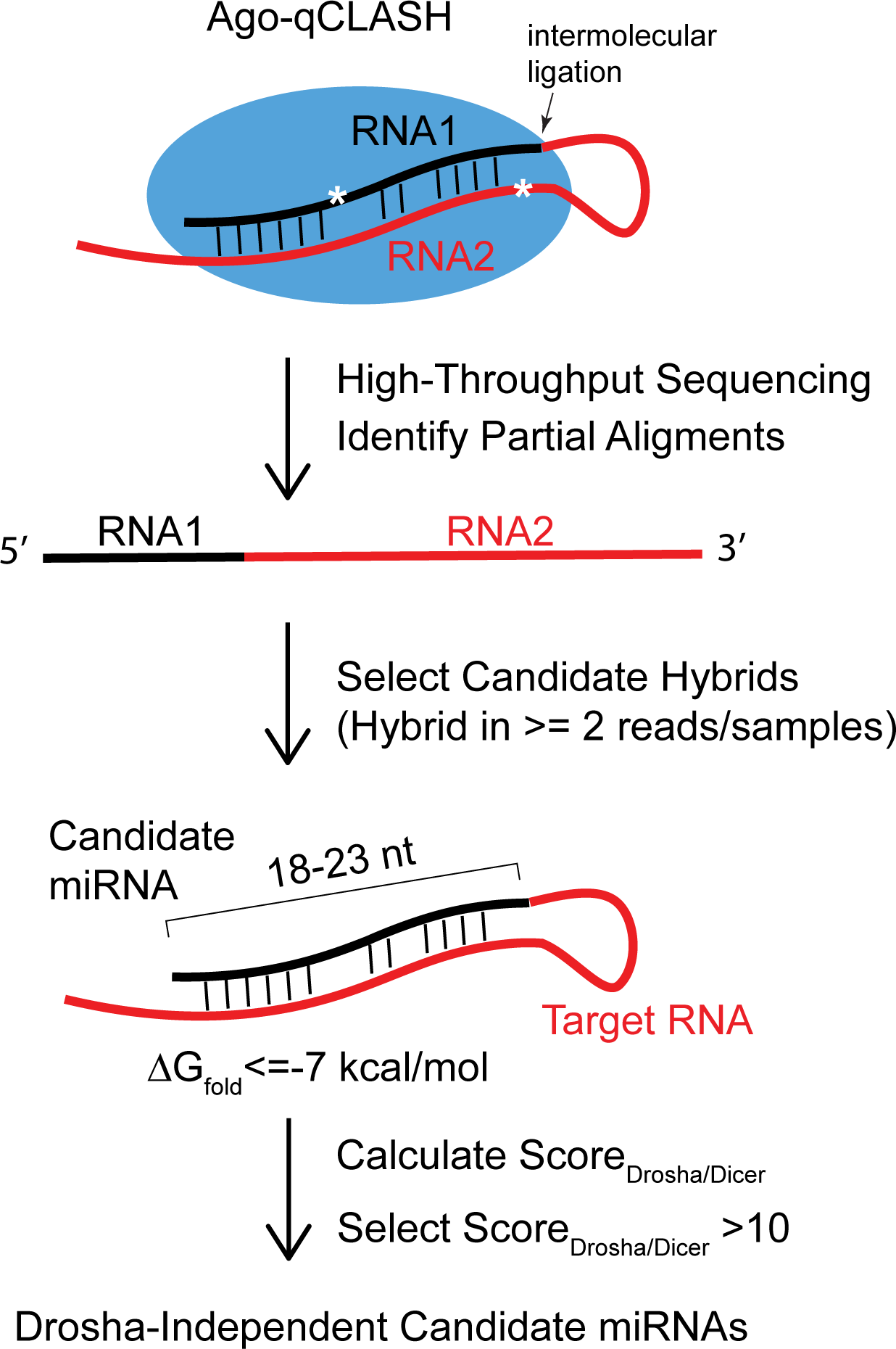
Target-Oriented microRNA Discovery (TOMiD) workflow. Following Ago-qCLASH, reads with partial alignments are evaluated as hybrids based on counts and biochemical parameters. Enrichment / suppression of candidate miRNAs is then calculated by comparison of the normalized counts in Drosha/Dicer conditions to wildtype, and the combined Score_Drosha/Dicer_ is calculated (See details in Supplemental Methods). * crosslinking between RNA and protein.

We applied the TOMiD method to detect Drosha-independent miRNAs in Ago-qCLASH data obtained from WT, Drosha-KO, and Dicer-KO HCT116 cells (Fields and Xie, Manuscript in preparation). Comparison between these datasets provides a unique platform to identify previously uncharacterized Drosha-independent miRNAs, which should be enriched in Drosha-KO cells but depleted in Dicer-KO cells. In total, we analyzed 13 Ago-qCLASH datasets, including 5 from WT cells, 4 from Drosha-KO cells, and 4 from Dicer-KO cells. To reduce identification of results arising due to sequencing error, each predicted unique cmiRNA sequence was required to exist in at least two non-identical hybrids per sample and be detected in two or more samples. To avoid the potential addition of bias to our analyses by clustering of predicted sequences, each cmiRNA was analyzed independently. This resulted in prediction of many highly similar cmiRNAs that map to the same RefSeq transcript but which vary by a few nucleotides in length or identity. As the final step in cmiRNA selection, we filtered cmiRNA that did not fall into a length range of 18-23 nt and which did not meet a predicted intra-hybrid folding energy (Δ*G*_*fold*_) of Δ*G*_*fold*_ ≤ −7.0 *kcal/mol* to specifically select for hybrids with highly miRNA-like characteristics based on miRNA/target sequence complementarity.

Over the 13 evaluated datasets, this process produced 106,021 unique cmiRNA sequences that aligned to 12,600 RefSeq transcripts, with many instances where several nearly-identical cmiRNAs were assigned to the same reference transcript. After initial alignment, several instances were identified in which cmiRNAs aligned to different RefSeq transcript variants arising from the same gene. For accurate quantification of results, in each of these cases a single reference transcript was selected as representative of that respective gene. A per-transcript combined count was then calculated by addition of all unique cmiRNA counts aligning to each respective RefSeq transcript. To account for potential differences in total cmiRNA counts across samples, proportional transcript counts (*C*_*cmiRNA*_) were calculated by dividing each combined cmiRNA count by the total counts per sample and a filter of an average *C*_*cmiRNA*_ ≥ 1.0 ∗ 10^+,^ across samples was applied (representing ∼25 average raw hybrid counts per transcript per sample).

Based on the results of previous studies, we hypothesized that Drosha-independent miRNAs would be identifiable in hybrid reads from WT samples, enriched in the Drosha-KO samples, and depleted in the Dicer-KO samples. To encompass all of these characteristics, we developed a scoring metric (*score*_*Drosha/Dicer*_) that combines cmiRNA enrichment in Drosha-KO cells and depletion in Dicer-KO cells compared to the average counts in the WT cells (details available in the Supplementary Methods). After hybrid prediction, filtration, and alignment of cmiRNAs to transcripts, we applied this scoring metric to the per-transcript combined cmiRNA counts to identify Drosha-independent cmiRNAs based on a high *score*_*Drosha/Dicer*_(range 0 to 501).

To select for high-confidence Drosha-independent miRNA candidates, we chose a threshold of *score*_*Drosha/Dicer*_ ≥ 20 which identified six high-confidence cmiRNAs for further analysis (Table 1). Of these, five cmiRNA were identified as known Drosha-independent miRNAs: miR-886 (vtRNA2-1), miR-3615, miR-877, miR484, and miR-320 (Berezikov et al. 2007; Babiarz et al. 2008; Persson et al. 2009; Kim et al. 2016). The other top-scoring cmiRNA was not identified as a previously known miRNA gene, and instead was identified as arising from the small NF90-Associated RNA A (snaR-A) family of transcripts (Figs. 2A and 2B). The snaR-A transcripts are ∼120 nt, highly structured RNAs that interact with NF90 via a double-stranded RNA binding motif. The human genome contains 14 nearly-identical copies of snaR-A located on chromosome 19 (Parrott et al. 2011), which are transcribed by RNA Polymerase III (Pol III) by an intragenic promoter, with sequences of different members of the snaR-A family varying at only two nucleotide positions (Fig. 2A) (Parrott and Mathews 2007; Parrott et al. 2011). While a previous bioinformatic study suggested that a Dicer-dependent small RNA may derive from snaR-A, it did not identify any relationship to Ago that might indicate miRNA function (Langenberger et al. 2013).

**Figure 2.**
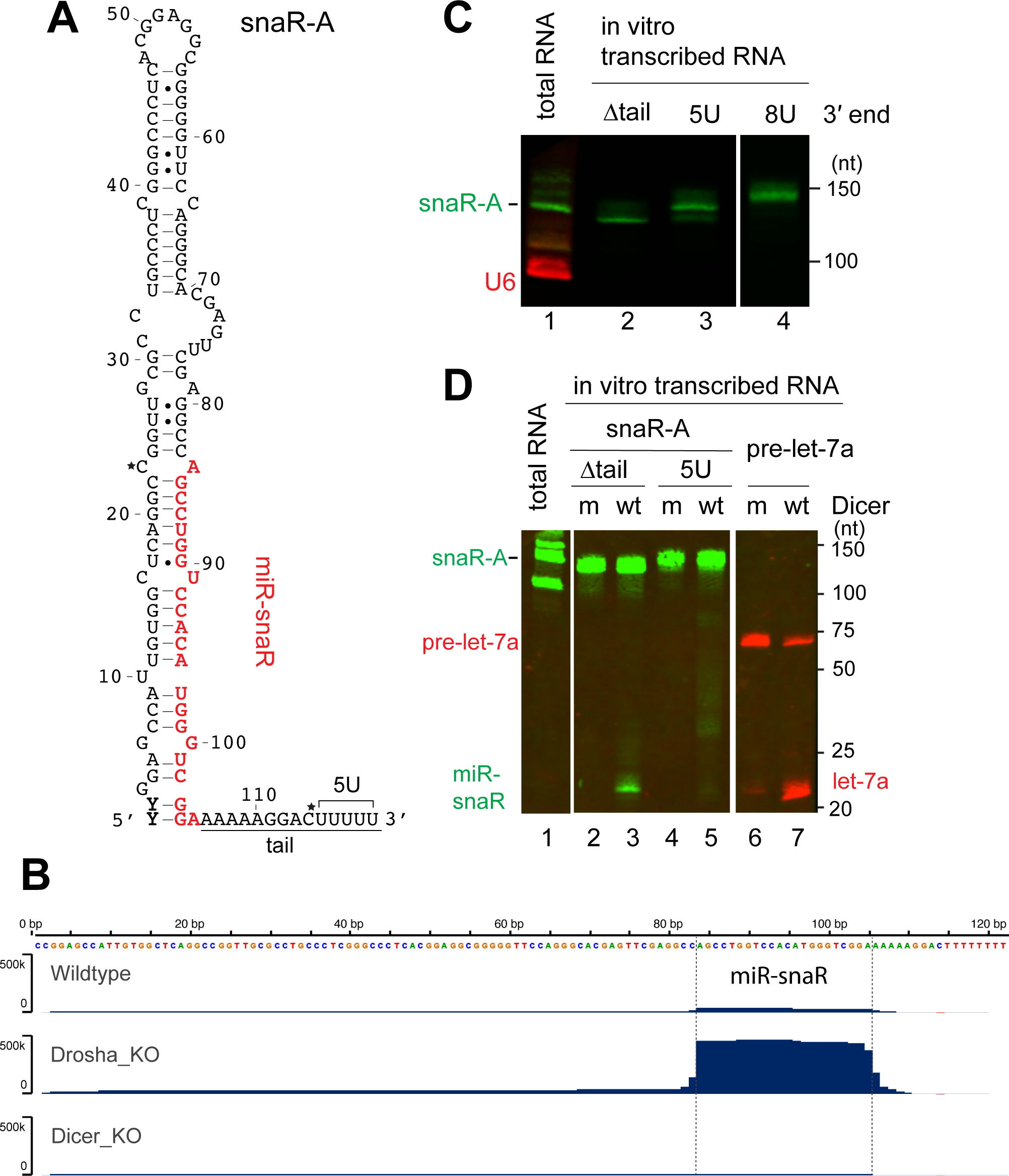
miR-snaR can be released from a truncated snaR-A transcript by Dicer. (**A**) Predicted secondary structure of snaR-A1, with the miR-snaR sequence highlighted in red. The two nucleotides that are variable within the 14 snaR-A family members are marked by an asterisk (*). (**B**) Histogram showing small RNA reads mapped to the snaR-A1 locus. Reads are identified in Ago-qCLASH experiments from HCT116 WT, Drosha-KO, and Dicer-KO cells. The boundaries of the miR-snaR are indicated by dotted lines. (**C**) Northern blot detection of various *in vitro* transcribed snaR-A transcripts, as well as endogenous snaR-A and U6 in total RNA extracted from 293T cells. (nt: nucleotides) (**D**) *In vitro* Dicer processing assay. *In vitro* transcribed pre-let-7a and snaR-A with or without the tail [illustrated in (A)] were processed by purified flag-tagged Dicer and analyzed by northern blot (wt: wildtype Dicer.; m: Dicer with mutated RNase III domains).

**Table 1.** Top-scoring Drosha-independent miRNAs identified by TOMiD. Column 2: RefSeq Transcript Identifier of aligned cmiRNA(s); Column 3: Consensus cmiRNA location in transcript; Column 4: human miRNA name or gene name (human miRNA “hsa-“ prefix omitted). (*) miR-886 gene symbol has been withdrawn by HGNC; Columns 5-7: Average hybrid counts per million (CPM) for each miRNA for WT (5 samples), Drosha-KO (4 samples), and Dicer-KO (4-samples), (counts per million candidate hybrids identified by TOMiD); Column 8: Enrichment score in Drosha-KO vs Dicer-KO cells (see Supplemental Methods).

| # | RefSeq | Consensus | Gene/miRN | CPM | CPM | CPM | Score<sub>Drosha/Dicer</sub> |
| --- | --- | --- | --- | --- | --- | --- | --- |
|  | Transcript | cmiRNA nts. | A name | WT | Drosha-KO | Dicer-KO |  |
| 1 | NR_030583.3 | 1-23 | vtRNA 2-1/<br>miR-886 | 24.1 | 97.9 | 0.0 | 501.4 |
| 2 | NR_037409.1 | 51-73 | miR-3615 | 10.4 | 36.5 | 0.0 | 318.7 |
| 3 | NR_030615.1 | 1-22 | miR-877 | 57.6 | 294.8 | 1.0 | 281.2 |
| 4 | NR_030159.1 | 8-28 | miR-484 | 50.5 | 149.8 | 0.9 | 181.4 |
| 5 | NR_004435.1 | 84-105 | snaR-A | 37.7 | 738.7 | 5.9 | 132.2 |
| 6 | NR_029714.1 | 48-69 | miR-320a | 1907.4 | 3803.9 | 86.8 | 46.5 |

### Characterization of the snaR-A sequence

The potential miRNA derived from snaR-A, which we describe as miR-snaR, is located at the base of the snaR-A hairpin, followed by a ∼15 nt 3’ tail (Fig. 2A). To elucidate miR-snaR processing, we examined the endogenous snaR-A sequences as both 5’ and 3’ termini of a pre-miRNA are crucial elements for mature miRNA production (Ma et al. 2004; Park et al. 2011). We adapted rapid amplification of cDNA ends (RACE) to determine the 5’ and 3’ ends of the snaR-A endogenously expressed in human embryonic kidney (HEK) 293T cells. Sanger sequencing results reveal a stretch of at least 5 uridines (U) at the 3’ end of snaR-A, which acts as an efficient termination signal for Pol III transcription (Supplemental Fig. S1A). However, mixed nucleotide signals were detected from the 5’ end of the snaR-A (Supplemental Fig. S1B). We then cloned individual sequences from 5’ RACE products and found that snaR-A contain heterogenous sequences in the first two nucleotides at the 5’ end (Figs. 2A and Supplemental Fig. S1C). These findings are mostly consistent with previous reports, which also found heterogenous 5’ di-nucleotide, but indicated that as many as 8 Us could be present at the 3’ end of snaR-A (Parrott and Mathews 2007).

To further investigate the length of endogenous snaR-A transcripts, we *in vitro* transcribed three different versions of snaR-A, including either 5 or 8 consecutive Us as the 3’ terminal nucleotides (5U and 8U), as well as a truncated snaR-A (Δtail) whose 3’ end coincides with the potential miR-snaR. We reasoned that a truncated snaR-A is more likely to be the direct substrate for Dicer, because a 3’ tailed hairpin is not a typical Dicer substrate (Ma et al. 2004). We analyzed these three transcripts alongside with total RNA extracted from 293T cells on an irNorthern blot (Fig. 2C) (Miller et al. 2018; Fields et al. 2019). The major population of snaR-A from 293T cells migrate at the same position as the snaR-A containing 5Us (Fig. 2C, compare lanes 1 and 3). Therefore, both our RACE and northern blot data suggest that the majority of endogenous snaR-A transcripts have 5 Us at the 3’ end.

### miR-snaR biogenesis is independent of Drosha, but depends on XPO5 and Dicer

As mentioned before, the full length snaR-A is unlikely to be processed directly by Dicer due to the long 3’ tail, while snaR-A Δtail resembles a traditional pre-miRNA with short 3’ overhang. Indeed, when we subject both snaR-A full length and Δtail to FLAG-tagged Dicer purified from HEK 293T cells, only the snaR-A Δtail was successfully processed into miR-snaR (Fig. 2D, compare lanes 3 and 5). However, no such snaR-A Δtail intermediate is abundantly detected in the total RNA sample (Fig. 2C, compare lanes 1 and 2; Fig. 2D, compare lanes 1 and 2). It is also unclear how the tail region is removed from the full length snaR-A for subsequent Dicer processing. Nonetheless, a hairpin without the 3’ tail would be compatible with both XPO5-mediated export and subsequent Dicer cleavage, and our Dicer processing assays validate that the potential miR-snaR can indeed originate from snaR-A.

Next, we explored the biogenesis of miR-snaR in HCT116 cells, in which snaR-A is not highly expressed (Supplemental Fig. S2A). We cloned the endogenous snaR-A gene into the pBlueScript plasmid and transfected the plasmid into HCT116 cells to examine miR-snaR expression. In three additional HCT116 cell lines we used for transfection, essential genes for canonical miRNA biogenesis were knocked out by CRISPR-Cas9, including Drosha, Dicer and XPO5 (Kim et al. 2016). Expression of miR-snaR in these HCT116 cells was analyzed by northern blot after Ago-immunoprecipitation (IP). MiR-snaR is clearly detectable in the WT HCT116 cells (Fig. 3A, lanes 1-3), again confirming that miR-snaR can be generated from the full-length snaR-A transcript. Compared to WT HCT116 cells, knockout of Drosha resulted in an obvious increase of miR-snaR (Fig. 3A, compare lanes 3 and 6). In contrast, miR-snaR could not be detected in Dicer-KO cells, and is detectable at a lower level in XPO5-KO cells (Fig. 3A, lanes 9 and 12). In the small RNA-seq data from the these HCT116 cell lines, we also confirmed that the expression levels of endogenous miR-snaR are the highest in Drosha-KO cells, but diminished in Dicer-KO and XPO5-KO cells (Supplemental Fig. S2B) (Kim et al. 2016). Combined with results from the Dicer processing assay, we conclude that miR-snaR is a non-canonical miRNA dependent on Dicer and XPO5 but independent of Drosha.

**Figure 3.**
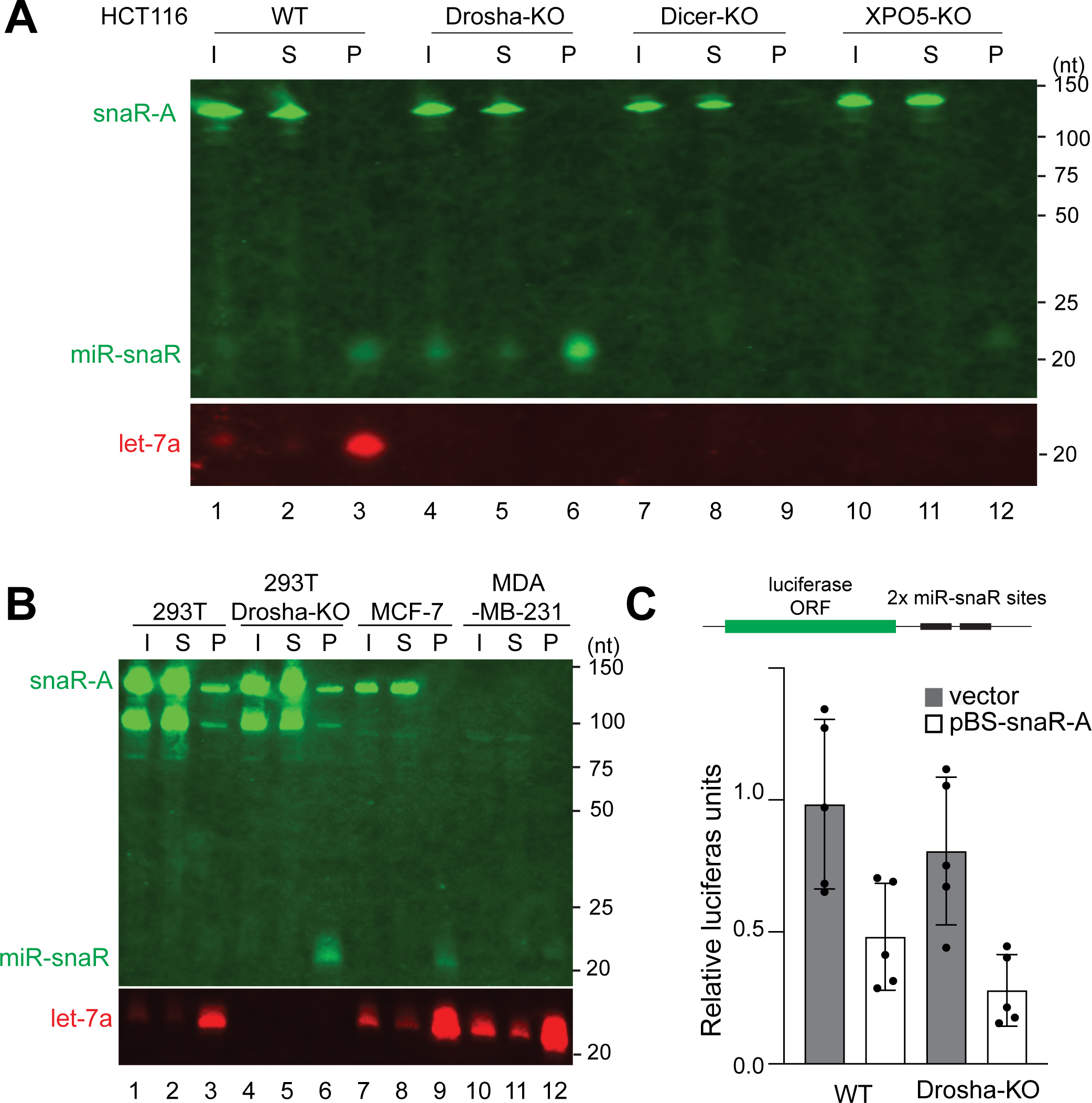
Functional miR-snaR associates with Ago in cells. (**A**) Ago-IP was performed with total cell lysate extracted from HCT116 cells (WT and knockouts) transfected with pBS-snaR-A. RNAs associated with Ago were extracted and analyzed by northern blot to detect snaR-A, miR-snaR, and let-7a. (**B**) Ago-IP and northern blot were performed as in (A) with samples from 293T, MCF-7, and MDA-MB-231 cells. Input (I) and supernatant (S) are 5% relative to the pellet (P) (nt: nucleotides). (**C**) Top: dual luciferase reporter for miR-snaR (pmirGLO-snaR), in which target site complementary to miR-snaR is cloned downstream of firefly luciferase, was transfected in 293T cells together with either pBS-snaR-A or pBS. Bottom: firefly luciferase activity was measured 48 hr post transfection and normalized to Renilla luciferase activity. Error bars represent standard deviation from five experiments. All differences are statistically significant (Student’s *t*-test, p<0.05).

### Detection of miR-snaR and its repressive function in cells

To examine if functional miR-snaR is expressed endogenously in cells, we performed Ago IP in HEK 293T and breast cancer cell lines that are known to express snaR-A (Parrott and Mathews 2007; Lee et al. 2016; Lee et al. 2017), which may lead to a higher endogenous level of miR-snaR. RNAs co-immunoprecipitated with Ago were analyzed by northern blot (Fig. 3B). While miR-snaR was not readily detectable in HEK293T cells, knocking out Drosha dramatically enriched miR-snaR (Fig. 3B, compare lanes 3 and 6), consistent with the results of our previous analysis (Table 1 and Fig.3A). In HER2 positive breast cancer MCF-7 cells, endogenous miR-snaR are detectable after Ago-IP (Fig. 3B, lane 9). Interestingly, the levels of miR-snaR and snaR-A in MCF-7 and 293T cells do not correlate, indicating that cell-specific factors contribute to the efficiency of miR-snaR biogenesis (Fig. 3B, compare lanes 1-3 to lanes 7-9). In triple negative breast cancer MDA-MB-231 cells, neither snaR-A nor miR-snaR was detected by northern blot (Fig. 3B, lanes 10-12), contradicting a previous report of high snaR-A expression in this cell line (Lee et al. 2017). In conclusion, we confirm that miR-snaR is endogenously expressed in cell lines that express snaR-A.

To determine if mature miR-snaR can function in RNA interference, we performed luciferase reporter assay in both WT and Drosha-KO HEK293T cells. Cells were transfected with either a plasmid expressing full-length snaR-A or an empty vector. Additionally, cells were co-transfected with a luciferase reporter plasmid with a miR-snaR binding site consisting of two repeats of a sequence that is fully complementary to miR-snaR in the 3’UTR of the luciferase gene. Luciferase signal was then measured 48 hours after transfection. In both WT and Drosha-KO cells the luciferase signal in groups expressing snaR-A was reduced approximately 50-60 % compared to their respective controls (Fig.3C). Of note, the luciferase signal in the Drosha-KO control was lower than that in the WT control. As a Drosha-independent miRNA, endogenous miR-snaR, as well as other Drosha-independent miRNAs, is expected to be elevated in Drosha-KO cells (Table 1 and Fig. 3A), which accounts for the comparatively lower luciferase signal. The result of this assay confirms that miR-snaR is processed from the full-length snaR-A and that the resulting miR-snaR is biologically active. These results fully complement and support the results of previous experiments examining the biogenesis and Ago-interaction of miR-snaR (Figs. 3A and 3B).

### Identification of miR-snaR targets in Ago-CLASH datasets

A number of previous studies have revealed snaR-A’s involvement in metabolic dysregulation of breast, liver, and ovarian cancers (Lee et al. 2016; Lee et al. 2017; Huang et al. 2018; Shi et al. 2019) and in drug-resistance in human colon cancer cells (Lee et al. 2014). We questioned whether miR-snaR can contribute to the phenotypes associated with snaR-A. To investigate this, mRNA targets of miR-snaR were identified from multiple Ago-CLASH and qCLASH datasets obtained from HCT116 WT and Drosha-KO cells, as well as from HEK293 cells (Helwak et al. 2013), by the Hyb software package using a custom reference database containing miR-snaR.

We first used UNAFold to analyze the pattern of miR-snaR/target interaction (Markham and Zuker 2008a; Travis et al. 2014). The hybkit software package (version 0.2.1a) developed by the Renne Lab was utilized to select hybrids containing mRNA and mRNA-pseudogenes and the frequency of pairing interaction at each nucleotide position of miR-snaR was calculated (Fig. 4A, left graph). Previous studies utilizing Ago-CLASH have observed hybrids with a higher contribution of significant or exclusive non-seed miRNA/target interactions than expected based on canonical understanding of seed-focused miRNA targeting. The observed targeting interactions of miR-snaR conversely display primarily seed-focused pairing. Within miR-snaR hybrids, an average of 84.0% of interactions include base paring of the seed region (nts 2-8). However, a region of significant supplemental base-pairing is also observed towards the 3’ end of miR-snaR (nts 15-17, average 88.2%) (Fig. 4A, left graph). When considering the entire miRNA/target dataset (excluding snaR-miR), a relatively even distribution of target interaction across the complete miRNA length was observed (Fig. 4A, right graph), consistent with previous observations by Gay et al. (Gay et al. 2018). The seed-focused interactions of miR-snaR suggest a potentially high potency in mediating target gene repression that is observed with canonical seed-focused miRNA/target interactions (Bartel 2018).

**Figure 4.**
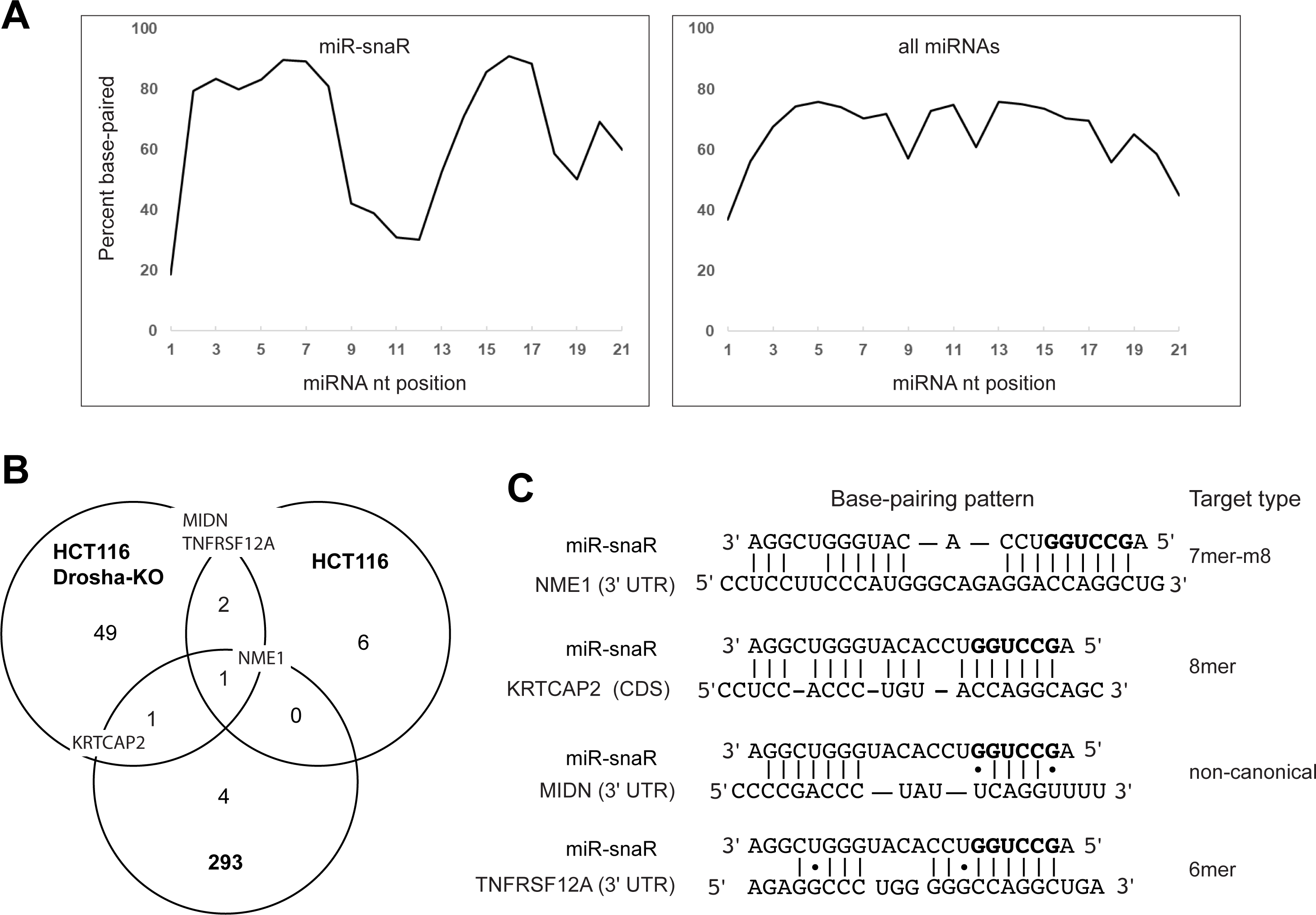
Identification of miR-snaR targets. (**A**) The intra-hybrid base pairing pattern between the miRNA and its target mRNAs are predicted by UNAFold using Ago-qCLASH hybrid reads for miR-snaR (left) and all other miRNAs (right). (**B**) Venn diagram showing seed-matched miR-snaR targets that appear in two or more hybrids from three Ago-CLASH datasets: 293, HCT116 and HCT116 Drosha-KO. Targets that appear in multiple datasets are shown. (**C**) Base-pairing pattern between miR-snaR and targets identified in (B). Watson-Crick base pairs are represented by lines. G-U wobble base pairs are represented by dots.

We next examined specific miR-snaR targets identified in Ago-CLASH datasets in detail. Based on miR-snaR’s predominantly seed-focused binding, targets were filtered to only mRNA targets in which a full seed match was present and in which two nonidentical hybrids were identified. This resulted in 63 identified targets across the three datasets. Targets were then selected where they were identified in at least two datasets, which provided four high-confidence targets of miR-snaR for detailed study (Fig. 4B). Three of these targets, KRTCAP2, MIDN, and TNFRSF12A, appear in two datasets. The target site in KRTCAP2 falls within the coding sequence, an atypical location for miRNA binding (Fig. 4C). For MIDN and TNFRSF12A, the target sites are complementary with the miR-snaR seed, but MIDN target site contains G-U wobble base pairing (Fig. 4C). Among the four high-confidence targets, nucleoside diphosphate kinase 1/non-metastatic cells 1 (NME1) was the only target to appear in all three datasets (Fig. 4B). Additionally, the putative miR-snaR binding site located within the 3’ UTR of NME1 has the characteristics of a canonical miRNA binding site. This site is located just ∼50 nt downstream from the stop codon. In addition, NME1 includes a target sequence that is fully complementary to the miR-snaR seed region and which is also fully conserved among the higher-order primate species in which the snaR-A gene exists (Fig. 4C and Supplemental Fig. S3). Together, these details strongly suggest NME1 as a genuine target of miR-snaR.

Interestingly, NME1 was the first human gene to be identified as a suppressor of cancer metastasis (Steeg et al. 1988). In breast cancer, an inverse correlation between NME1 and metastatic potential has been well documented (Leone et al. 1993; Horak et al. 2007). Studies have also shown that breast cancer cells secrete NME1, and this extracellular NME1 supports primary tumor growth and metastasis (Yokdang et al. 2015). Given that snaR-A promotes proliferation and migration of breast cancer cells (Lee et al. 2017), miR-snaR may cooperate with snaR-A by repressing NME1 expression.

### miR-snaR reduces NME1 mRNA levels via a specific binding site in the 3’ UTR

To examine whether miR-snaR can negatively impact NME1 mRNA levels, we performed reverse transcription followed by real time PCR (RT-qPCR) after overexpressing miR-snaR in HEK 293T cells where miR-snaR is not highly expressed (Fig. 3B). Introduction of miR-snaR was performed either by plasmid transfection or by transfection of a synthetic miRNA mimic. (Figs. 5A and 5B). This experimental setup allows identification of functions of miR-snaR independent of the full length snaR-A transcript, which may influence gene expression through a different mechanism. The plasmid was designed to express a U6 promoter-driven short hairpin RNA (shRNA) to be subsequently cleaved by Dicer and produce miR-snaR (Fig 5A). As a control, a plasmid expressing a shRNA with a scramble sequence of miR-snaR hairpin was constructed. Northern blot confirmed the expression of miR-snaR from the utilized plasmid (Fig. 5A, northern blot). In both miR-snaR mimic and plasmid transfection experiments, NME1 mRNA is significantly reduced in cells concurrent to elevated levels of miR-snaR (Fig. 5C). The results of this experiment provide evidence that the NME1 mRNA can indeed be regulated by miR-snaR in cells.

**Figure 5.**
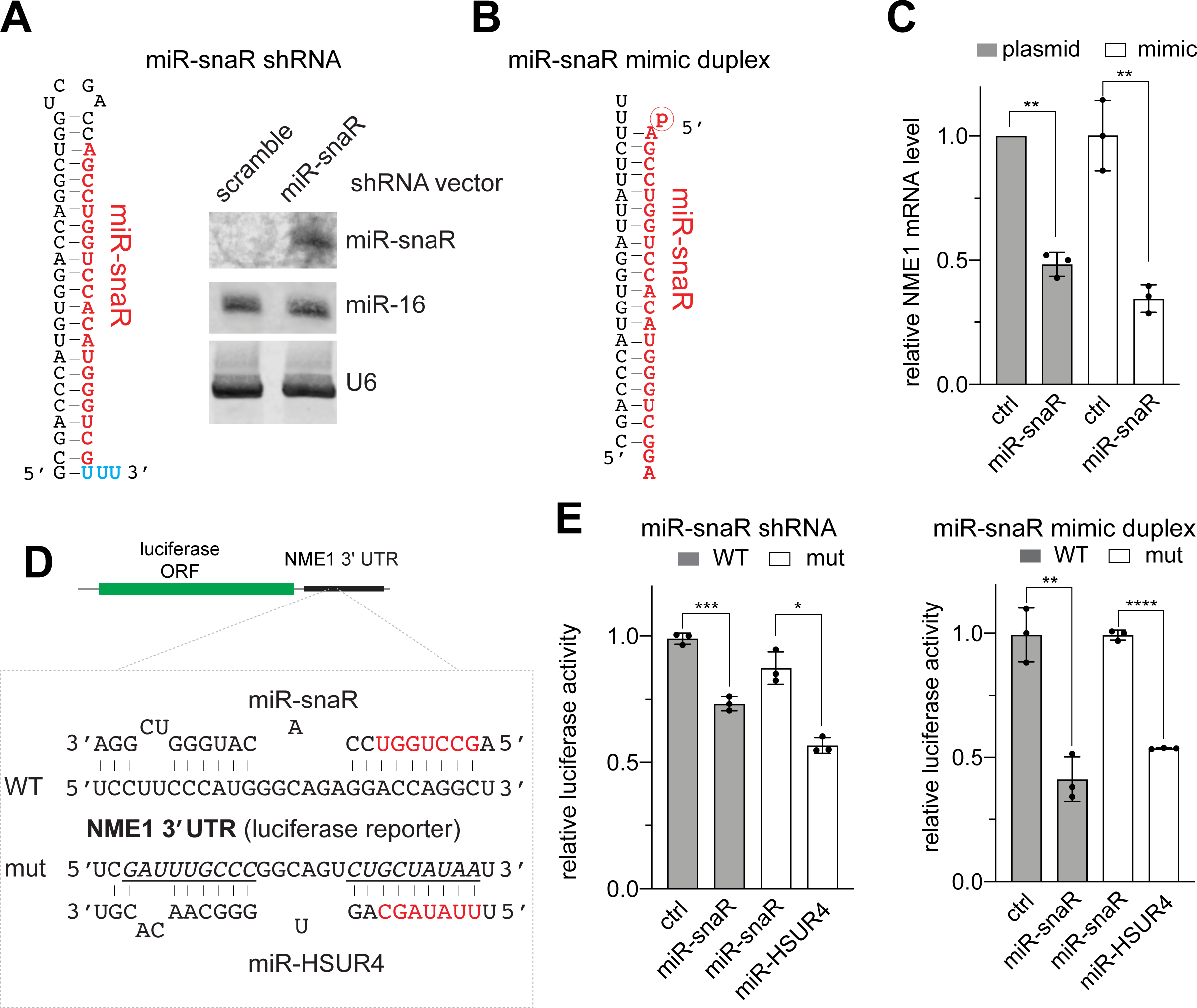
miR-snaR represses NME1 mRNA by interacting with its 3’ UTR. (**A**) Left, schematic of the shRNA encoded in pU6-miR-snaR, with miR-snaR sequence highlighted in red. The terminal Us (cyan) are required for efficient RNA Pol III termination. Right, northern blot analyzing the levels of miR-snaR in total RNA extracted from 293T cells transfected with either pU6-scramble-shRNA or pU6-miR-snaR. Endogenous miR-16 and U6 serve as internal loading controls. (**B**) Schematic of the miR-snaR mimic duplex. The miR-snaR mimic strand in red is modified with a 5’ phosphate group. (**C**) RT-qRCR analysis of NME1 mRNA in total RNA extracted from HEK293T cells transfected with pU6-miR-snaR or miR-snaR mimic. NME1 mRNA expression was normalized to β-Actin. Three biological replicates for each sample were recorded and data was graphed and analyzed using Prism GraphPad. **p≤0.01. (**D**) Depiction of WT and mutant NME1 target sites on the luciferase reporter along with their targeting miRNA: miR-snaR and miR-HSUR4, respectively. Seed sequences of the miRNAs are highlighted in red. Regions mutated in the mutant reporter are underlined. (**E**) Dual luciferase reporter containing the WT or mut NME1 3’ UTR downstream of firefly luciferase was transfected in 293T cells together with either pU6-miR-snaR (left) or miR-snaR mimic (right). Firefly luciferase activity was measured 48 hours post transfection and normalized to Renilla luciferase activity. Error bars represent standard deviation from three experiments. All differences are statistically significant (Student’s *t*-test, *p≤0.05; **p≤0.01; ***p≤0.001; ****p≤0.0001).

Binding site specificity is critical to the regulatory function of miRNAs. To investigate whether the miR-snaR binding site on NME1 is the sequence captured by Ago-CLASH (Fig. 4B), we performed a dual luciferase reporter assay, in which the firefly luciferase reporter contained the 3’ UTR of NME1 (Fig. 5D). As a control, we also constructed a luciferase reporter plasmid with the miR-snaR binding site in NME1 3’ UTR mutated to base-pair with miR-HSUR4 (mut). This miR-HSUR4 control was chosen due to its viral origin (*Herpesvirus saimiri*) and its lack of endogenous expression in HEK293T cells. A luciferase assay was conducted using both the previously described plasmid and mimic transfection methods for miR-snaR expression (Figs. 5A and 5B). In both scenarios, miR-snaR significantly decreased luciferase expression when co-expressed with WT NME1 3’ UTR reporter, but not when co-expressed with a mut NME1 3’ UTR reporter (Fig. 5E). In contrast, luciferase signal in cells transfected with mut NME1 3’ UTR reporter was reduced by miR-HSUR4, indicating that this miRNA binding site is responsible for target repression.

### miR-snaR downregulates NME1 protein to promote cancer cell migration

After confirming that miR-snaR reduces NME1 mRNA within cells, we then analyzed NME1 protein expression. We transfected two cell lines – HEK293T and MDA-MB-231– with miR-snaR mimic, as these cells have low levels of endogenous miR-snaR expression (Fig. 3B, lanes 3 and 12). Additionally, MCF-7 cells, in which endogenous miR-snaR is detectable, were transfected with miR-snaR inhibitor to reverse potential miR-snaR-mediated repression on NME1. In each case, total protein lysate was collected 48 hours after transfection and protein levels were determined by western blot. In both HEK293T and MDA-MB-231 cells, miR-snaR transfection resulted in a significant decrease in NME1 protein levels (Fig. 6A, lanes 1-4; Fig. 6B). On the other hand, inhibition of miR-snaR is observed to increase NME1 level in MCF-7 cells, albeit by a rather moderate amount (Fig. 6A, compare lanes 5 and 6; Fig. 6B). We also investigated NME1 protein levels in HCT116 cells, in which miR-snaR is upregulated in Drosha-KO cells but diminishes in Dicer-KO cells (Supplemental Fig. S2B). Supporting targeting of NME1 by miR-snaR, we observed an inverse correlation between NME1 and miR-snaR levels, with low levels of NME1 in Drosha-KO cells and high levels of NME1 in Dicer-KO cells, respectively (Fig. 6C, lanes 2 and 3). Importantly, introduction of miR-snaR inhibitor in Drosha-KO cells elevated NME1 (Fig. 6C, compare lanes 4 and 5), and introduction of miR-snaR mimic in Dicer-KO cells repressed NME1 (Fig. 6C, compare lane 6 and 7). These results suggest that NME1 protein levels are directly influenced by miR-snaR.

**Figure 6.**
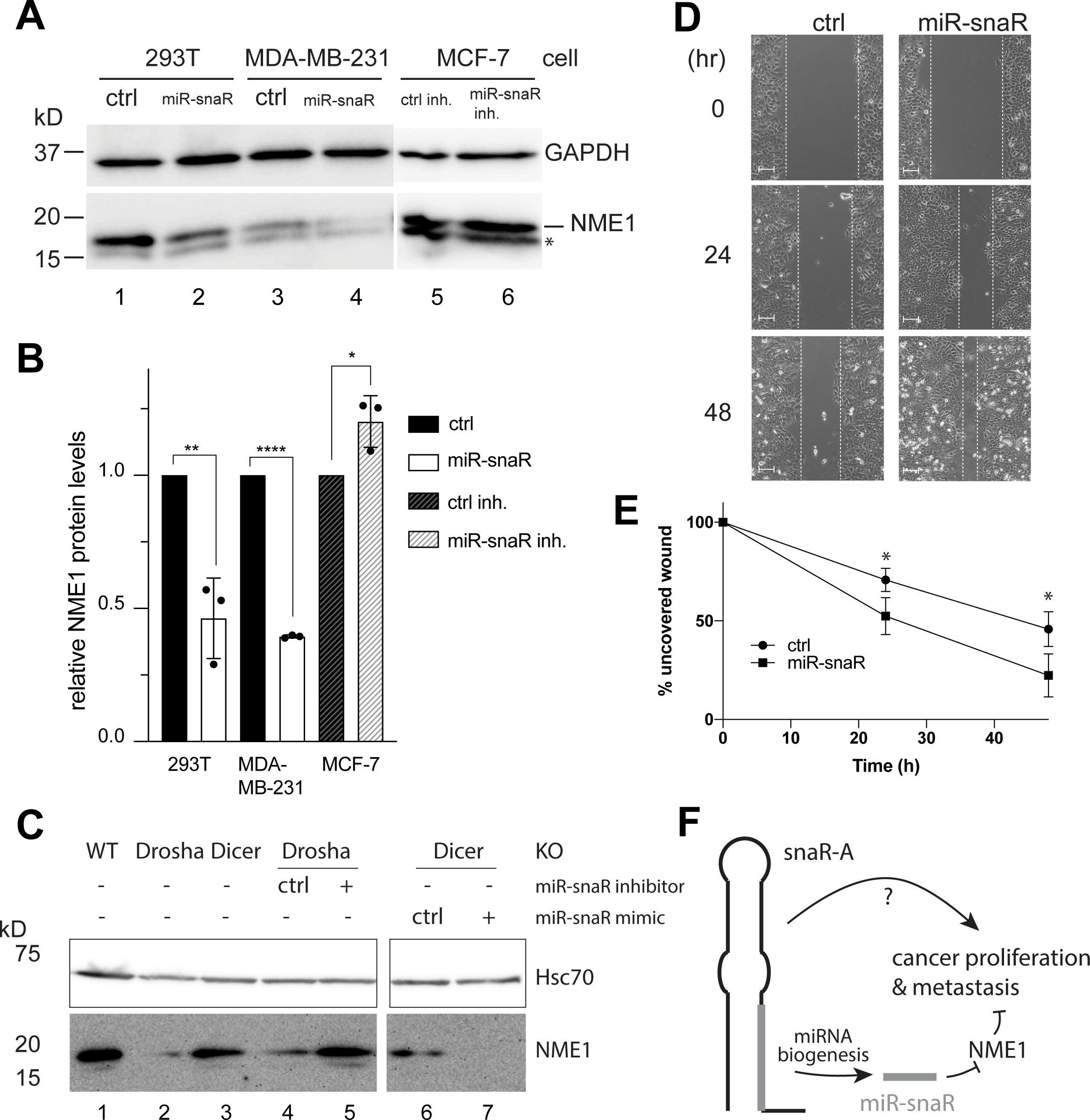
miR-snaR downregulates NME1 to promote cell migration. (**A**) Western blots show reduction of NME1 protein in 293T and MDA-MB-231 cells after transfection of miR-snaR mimic and increase of NME1 protein in MCF-7 cells after transfection of miR-snaR inhibitor, with GAPDH as a loading control. Asterisk (*) marks a non-specific band that is unlikely to be the reported smaller isoform of NME1, whose 3’ UTR also contains the miR-snaR target site. (**B**) Quantitation of three experiments in (A) is represented in a bar-graph. All differences are statistically significant (Student’s *t*-test, **p≤0.05; **p≤0.01; ****p≤0.0001). (**C**) Western blot of NME1 protein in HCT116 WT, Drosha-KO, and Dicer-KO cells. Drosha-KO and Dicer-KO cells were additionally transfected with miR-snaR inhibitor/control and miR-snaR mimic/control, respectively. Hsc70 is used as a loading control. (**D**) Wound healing assay of MCF-7 cells transfected with miR-snaR mimic or a control mimic. White lines outline the edge of the scratched wound. Scale bar: 100 µm. (**E**) Quantification of three experiments as shown in (D). (Student’s *t*-test, *p≤0.05). (**F**) Model of snaR-A’s role in enhancing tumor growth and metastasis. Derived from snaR-A, miR-snaR is inhibiting NME1 to promote metastasis.

Previous studies have shown that knockdown of snaR-A could inhibit proliferation and migration of breast cancer cells (Lee et al. 2016; Lee et al. 2017). Given this information, we looked further into the effect of miR-snaR in breast cancer cell lines. We utilized the MTT (3-(4,5-dimethylthiazol-2-yl)-2,5-diphenyltetrazolium bromide) assay to assess the effects of miR-snaR on cell proliferation. Considering that snaR-transcript knockdown inhibits proliferation (Lee et al. 2016; Lee et al. 2017), we speculated that increasing miR-snaR expression in MDA-MB-231 or MCF-7 cells would result in increased proliferative activity. To this end, MDA-MB-231 and MCF-7 cells were transfected with miR-snaR mimic. Additionally, HEK293T cells were also transfected. MTT assays were performed on cells 24 hours post transfection and immunoblotting was used to show reduction in NME1 protein as an indicator of successful transfection (Supplemental Figs. S4A and S4B). In contrast to our predictions, cell proliferation was neither reduced nor enhanced for any of the cell lines tested (Supplemental Fig. S4C).

As NME1 has a well-established role in inhibiting metastasis but not proliferation of cancer cells (Boissan et al. 2010; Marino et al. 2013), we reason that miR-snaR may specifically regulate cell migration. Potentially supporting this theory, it has been shown that MCF-7 cells have higher levels of NME1 and that siRNA-knockdown of NME1 increases cell mobility (Bemis and Schedin 2000). To examine this potential relationship, we performed a wound healing assay using MCF-7 cells transfected with either a control or a miR-snaR mimic. Consistent with our initial result, miR-snaR transfection in MCF-7 cells significantly reduced NME1 protein levels (Supplemental Fig. S4B). We also observed a significant increase in wound closure for miR-snaR-transfected cells over the course of 48 hours (Figs. 6D and 6E). Therefore, our data suggest that miR-snaR can promote cell migration by repression of NME1. Given the role of NME1 as a metastasis suppressor, our findings suggest an oncogenic role for miR-snaR in cancer.

## DISCUSSION

In this study, we developed the TOMiD method, an innovative bioinformatic approach for identification of novel miRNAs using the Ago (q)CLASH ribonomics technique (Fig. 1). Utilizing qCLASH datasets obtained from a panel of HCT116 cells lacking essential miRNA biogenesis factors (Drosha or Dicer), we discovered a miRNA (miR-snaR) that originates from snaR-A, a small non-coding RNA implicated in tumor proliferation and metastasis (Fig. 2). Despite being independent of Drosha processing, miR-snaR biogenesis is XPO5-and Dicer-dependent (Figs. 2 and 3). We showed that miR-snaR is biologically active due to its association with Ago and its ability to repress luciferase-reporter-containing complementary target sites (Fig. 3). Functionally, miR-snaR inhibits NME1 expression both at the mRNA and the protein levels (Figs. 4-6). Because NME1 is a metastasis inhibitor, we propose that snaR-A exerts its metastasis-promoting function at least in part through miR-snaR targeting of NME1 (Fig. 6F).

### TOMiD analysis for *de novo* miRNA discovery

The TOMiD approach expands the scope of Ago-(q)CLASH by enabling discovery of novel miRNAs in addition to the primary use of this technique for unambiguous identification of miRNA targeting interactions (Helwak et al. 2013). As demonstrated by the identification of miR-snaR, this methodology has a significant potential for use in characterization of previously unidentified noncanonical human miRNAs. In addition, TOMiD expands the ability of (q)CLASH to enable *de novo* miRNA characterization in non-model organisms or miRNA-producing pathogens, as it does not rely on preexisting annotations of miRNAs and requires only a reference transcriptome for miRNA discovery.

The TOMiD method also has a significant potential for further refinement and development to increase specificity in identification of miRNAs. In the current analysis, a low “signal-to-noise” ratio was observed with many non-miRNA hybrids predicted for each valid miRNA interaction, albeit with a characteristically lower count for occurrences in nonspecific predictions. While differentiation of potentially important novel or previously-known miRNAs can be made based on hybrid counts, performing a detailed characterization of the cutoff parameters utilized for filtration at different stages in the method and the addition of further data preprocessing steps each has the potential to significantly increase the specificity of miRNA-containing hybrid prediction and thus increase utility in novel miRNA discovery. Additionally, this method has the potential for future modification to utilize genomic, rather than transcriptomic reference sequences, allowing characterization of small sequence fragments that may escape detection as valid transcripts and facilitating the study of non-model organisms without a robust reference transcriptome. This method is currently under further development and is being implemented as a publicly-available analysis pipeline to be described elsewhere.

### MiR-snaR biogenesis and function

Previously, a small RNA sequence derived from the snaR-A transcript in a Dicer-dependent fashion was identified via a retrospective data analysis (Langenberger et al. 2013). However, whether this sequence could be classified as a miRNA was debatable. It has been shown that snaR-A is predominantly distributed in the cytoplasm of HEK293T and HeLa cells (Parrott et al. 2011). This cytoplasmic distribution of snaR-A is consistent with its accessibility to Dicer and other RNA interference machineries. Our discovery that miR-snaR associates with Ago and the ability of miR-snaR to suppress NME1 argues that this sequence is a bona-fide miRNA. SnaR-A is a non-coding RNA that is most highly expressed in testis, followed by brain and placenta, and is specific to higher-order primates (Parrott and Mathews 2007). Additionally, the full snaR-A transcript is upregulated in many cancer cell lines (Parrott and Mathews 2007; Lee et al. 2016; Huang et al. 2018; Shi et al. 2019). While it would seem reasonable for miR-snaR expression to follow the expression pattern of snaR-A, we have observed instances of disproportional snaR-A and miR-snaR expression in different cell lines (Fig. 3B), suggesting that cell-specific factors can control the processing from snaR-A to miR-snaR. One such factor could be the enzyme(s) that removes the tail of snaR-A, before Dicer could process the hairpin (Figs. 2A and 2D). The identity of this enzyme(s) requires further investigation. MiR-snaR can potentially arise by processing of any of the fourteen members of the snaR-A family, as each homolog has an identical sequence in the miR-snaR region (Fig. 2A). This abundant genetic source may explain the large degree of miR-snaR up-regulation in Drosha-KO cells (Table 1 and Supplemental Fig. S2B). In light of this, miR-snaR levels may be particularly sensitive to perturbation of Drosha levels. Indeed, in HCT116 Drosha-KO cells, miR-snaR rises to the second most abundant miRNA (Supplemental Fig. S2B and data not shown). Therefore, miR-snaR may be most influential in cells with low Drosha expression.

The discovery of miR-snaR adds to the ever-growing list of miRNAs that are derived from other small non-coding RNAs (Babiarz et al. 2008; Ender et al. 2008; Bartel 2018; Lemus-Diaz et al. 2020). Compared to canonical pre-miRNAs that are largely consumed to produce mature miRNAs, small non-coding RNAs are less efficiently processed into miRNAs, presumably because they are shielded by specific RNA-binding proteins. Also, small non-coding RNAs usually deviate from their normal conformation to adopt a hairpin that may not be optimal for Dicer processing (Babiarz et al. 2008; Lemus-Diaz et al. 2020). For example, Dicer cleavage efficiency of snaR-A hairpin is considerably lower compared to the pre-let-7a hairpin (Fig. 2D, compare lanes 3 to 7). Consequently, miRNAs originating from small non-coding RNAs are generally less abundant compared to most canonical miRNAs. Nonetheless, miR-snaR occurs in greater abundance than its peers as discussed above. These observations suggest that miR-snaR plays a substantial role in gene regulation.

In miR-snaR/target RNA hybrids obtained from Ago-(q)CLASH datasets, we observed seed-focused interactions (Fig. 4A, left) suggesting that miR-snaR is predominantly involved in canonical seed-match target-regulation. Based on identification of four miR-snaR targets found in more than two qCLASH datasets each, we predicted four genes (NME1, KRTCAP2, MIDN, and TNFRSF12A) as high-confidence targets of miR-snaR. Among these, miR-snaR targeting of NME1 is particularly interesting. First, the miR-snaR target site in NME1 is located only 56 nt downstream of the stop codon, a preferred site for miRNA-mediated repression in the 3’ UTR (Grimson et al. 2007), with absolute conservation among the higher-order primate species in which the snaR-A gene exists (Supplemental Fig. S3). The combination of the functional target location with this conservation suggests a functional importance for this targeting. Second, NME1 has been shown both to regulate expression of genes associated with metastasis and to limit cell motility through interactions with proteins such as Gelsolin (Horak et al. 2007; Marino et al. 2013; McCorkle et al. 2014). Combined with the role of NME1 as a well-known metastasis inhibitor and the demonstrated role of snaR-A in the promotion of metastasis in a variety of cancers, our identification of NME1 targeting by miR-snaR suggests that these two ncRNAs may play complementary roles in promoting cancer cell growth and/or migration. Furthermore, other than miR-snaR, two potential PIWI-interacting (pi-)RNAs can derive from snaR-A (Parrott and Mathews 2007). Given that the highest expression of snaR-A occurs in the testis, the existence of potential piRNA fragments derived from snaR-A is intriguing. Additionally, one of the piRNAs, the 27 nt long piR-30611, covers the entire miR-snaR sequence (Parrott and Mathews 2007). Whether and how these piRNAs may coordinate with miR-snaR and snaR-A in gene regulation warrants further investigation.

## MATERIAL AND METHODS

### Bioinformatic Analysis

To identify novel miRNAs, the TOMiD analysis method was implemented as a series of stepwise analyses (see Supplemental Methods for full implementation details). Sequence reads from Ago-qCLASH experiments (GEO accession number: GSE164634) in HCT116 cells were first analyzed for sequence quality and length utilizing FastQC. Sequencing adapters were removed by Trimmomatic, and Illumina read mate pairs were merged using PEAR (Zhang et al. 2014). PCR-duplicates were then removed using fastx-collapser, followed by removal of individual read barcodes by cutadapt. Reads were aligned to the hOH7 reference genome supplied with the Hyb software package using Bowtie2 (Langmead and Salzberg 2012; Travis et al. 2014), and the longest partial alignment was identified for each read. Aligned reads were then evaluated as potential hybrid sequences using a custom script implemented in Python3 using the pysam module with Samtools (Li et al. 2009). Several properties were then investigated for each hybrid. The Gibbs free energy of hybrid formation was first predicted using UNAFold hybrid-min (Markham and Zuker 2008b) and candidate hybrids were filtered to require a free energy <= -7.0 kcal/mol. Each read was then split in to two portions, with the longer alignment selected as the “target” and the shorter portion as the candidate miRNA (cmiRNA). Hybrids were then retained where the cmiRNA was 18 to 23 bases in length and was represented in two or more candidate hybrids occurring in at least two independent Ago-qCLASH sequencing libraries.

For identification of possible Drosha-independent miRNAs, the counts of each cmiRNA were compared in Ago-qCLASH datasets across WT, Drosha-KO, and Dicer-KO HCT116 cells. A scoring metric was applied to stratify cmiRNAs significantly enriched in Drosha-KO cells and depleted in Dicer-KO cells, and cmiRNAs with *score*_*Drosha/Dicer*_ ≥ 20 were identified as high-confidence Drosha-independent cmiRNAs. For identification of miR-snaR targets, a reference database containing miR-snaR was prepared and the Hyb pipeline was utilized to identify CLASH hybrids containing miR-snaR with its associated targets (Travis et al. 2014). The Vienna hybrid pairing representations produced by the UNAFold portion of the Hyb pipeline were then analyzed to plot the overall and miR-snaR-specific patterns of miRNA positional base pairing frequency utilizing the Hybkit toolkit. A detailed description of analysis methods, as well as the custom analysis scripts utilized, have been supplied in the Supplemental Materials.

### Plasmid construction

For expression of the snaR-A transcript, 555 base-pairs of genomic sequence encompassing the snaR-A sequence was cloned into the pBlueScript (pBS) plasmid using EcoRI and XhoI restriction enzyme sites and the resulting plasmid was named pBS-snaR-A. The cloned snaR-A with surrounding sequences is identical to either snaR-A4, 7, 8, 9, 10, or 11. To express miR-snaR independently of the full length snaR-A, we first cloned the U6 promoter into pBS between EcoRI and XhoI sites. We then assembled a short hairpin (Fig. 5A) through overlapping PCR and cloned it downstream of the U6 promoter by quick change mutagenesis to produce pU6-miR-snaR. The pU6-scramble-shRNA was constructed by replacing miR-snaR shRNA with a scrambled sequence also using quick change mutagenesis. To test miR-snaR’s gene silencing effect, pmirGLO-snaR was constructed on pmirGLO Dual-Luciferase miRNA target expression vector (Promega, E1330). Two oligonucleotides were annealed to form two repeats of a sequence that is reverse complementary of the miR-snaR sequence, which was then inserted into pmirGLO via XbaI and SacI restriction enzyme sites. To confirm the miR-snaR binding site in the 3’ UTR of NME1 mRNA, pmirGLO-NME1 and pmirGLO-NME1-mut were generated on pmirGLO vector. To construct pmirGLO-NME1, cDNA of the last exon of the NME1 mRNA including the 3’ UTR was inserted into pmirGLO via XbaI and SacI sites. The putative miR-snaR binding site was then mutated by site directed mutagenesis in pmirGLO-NME1-mut (Fig. 5C). All primers utilized in cloning are listed in Supplemental Table S1.

### *In vitro* transcription and Dicer processing assays

Polymerase chain reaction (PCR) DNA templates for snaR-A transcription were amplified from pBS-snaR-A with a forward primer containing the T7 promoter sequence and various reverse primers to generate different versions of snaR-A transcripts. These included a truncated snaR-A (Δtail), as well as 5 or 8 consecutive Us (5U and 8U) as the 3’ termination nucleotides. The 5’ variable dinucleotides of the transcripts are set to GC to allow efficient T7 transcription. Utilized primers are listed in Supplemental Table S1. The DNA products were separated on a 3% agarose gel and then extracted using gel extraction kit (ZYMO research). *In vitro* transcription with T7 RNA polymerase and *in vitro* Dicer cleavage assays were performed as previously described (Sheng et al. 2018).

### Cell Culture, transfection, total RNA and protein extraction

HEK293T, breast cancer MCF-7, and breast cancer MDA-MB-231 cells were cultured in Dulbecco’s Modified Eagle Medium (DMEM, HyClone SH30243.FS) with 10% fetal bovine serum (FBS, Gibco) and 1% penicillin/streptomycin (P/S) at 37°C with 5% CO2. HCT116 colorectal cancer cells were cultured in McCoy’s 5A medium (HyClone, SH30200.FS) with 10 % FBS, 1 % P/S at 37°C, with 5% CO2.

Plasmid DNA was transfected using polyethylenimine (PEI) (Polysciences Inc, 24765-1) or Lipofectamine 3000 (Life Tech, L3000015) in HEK293T or HCT116 cells seeded 24 hours prior to transfection, according to the manufacturer’s instructions. MiRNA mimic or inhibitor was transfected using Lipofectamine RNAiMAX transfection reagent (Life Tech, 13778150) in HEK293T, MCF-7, and MDA-MB-231 cells in suspension, according to the manufacturer’s instructions.

Total RNA was isolated using TRIzol reagent (Life Tech, 15596018) and treated with RQ1 DNase (Promega, M6101) 48 hours after transfection, according to manufacturer’s instructions. After DNase treatment, RNA was purified using phenol-chloroform-isoamyl alcohol (25:24:1) (PCA) followed by ethanol precipitation. To collect total protein, approximately 200 µL of NP-40 lysis buffer (50 mM Tris–HCl pH 7.5, 1% NP-40, 10% glycerol, 150 mM NaCl, 5 mM EDTA, and 0.5 mM PMSF) was added per million cells. Cell lysate were mixed with gentle agitation for 30 minutes at 4°C and supernatant was collected by centrifugation at 21,000 × g for 10 minutes at 4°C.

### 5’ and 3’ RACE of the endogenous snaR-A

Total RNA was extracted from HEK293T WT and Drosh-KO cells. The 3’ terminus of total RNAs were ligated with the irCLIP adaptor by T4 RNA Ligase 1 (NEB, M0437M). After ligation, the RNAs were annealed with a cDNA synthesis primer and cDNAs were generated by SuperScript™ IV Reverse Transcriptase (Invitrogen, 18090050). The cDNA/RNA duplexes were then purified by streptavidin beads (Invitrogen, 65601) and the RNAs hydrolyzed by NaOH. The cDNA was circularized by CircLigase™ II ssDNA Ligase (Epicentre, CL9025K). Next, cDNA of snaR-A was amplified from circular single-stranded DNA templates by PCR using primers (P6 tall, P3 tall) that anneal to the irCLIP adaptor and the gene specific reverse primers (snaR-A 5’, snaR-A 3’). The PCR amplicons were purified by Zymoclean™ Gel DNA Recovery Kit (ZYMO RESEARCH, D4008) and phosphorylated at the 5’ termini by T4 Polynucleotide Kinase (NEB, M0201L). The pBluescript plasmid was cut by EcoRV and dephosphorylated by Calf Intestinal Alkaline Phosphatase (CIP) (NEB, M0290L), then ligated with the phosphorylated PCR fragments by T4 DNA Ligase (NEB, M0202S). Inserts were analyzed by Sanger sequencing. All primers used are listed in Supplemental Table S1.

### Immunoprecipitation, western blot and antibodies

Ago-IP was performed using α-Ago antibody (clone 4F9) as previously described (Sheng et al. 2018). Antibodies for NME1 (ABClonal, A0259), Hsc70 (Santa Cruz, sc-7298), and GAPDH (Cell Signaling, 14C10) were used for western blots.

### RNA quantification

Northern blot analyses with near infrared dye-labeled probes were performed as previously described (Miller et al. 2018; Fields et al. 2019). The probes utilized are listed in Supplemental Table S1. For RT-qPCR, 1µg of total RNA was reverse transcribed to generate cDNA using iScript RT Supermix (Bio-Rad, 1708841). The reaction was completed using the following protocol: 5 minutes at 25°C (priming), 20 minutes at 46°C (reverse transcription), 1 minute at 95°C (RT inactivation). Resultant cDNA was used for quantitative PCR (qPCR) using SYBR GreenER qPCR SuperMix (Bio-Rad, 1725275) using the standard qPCR protocol after initial denaturing step at 95°C for 30s, repeating 40 cycles (95°C 15s, 60°C 30s) followed by melting curve analysis.

NME1 mRNA levels were normalized against β-Actin mRNA levels using 2^-ΔCt^. Primers utilized are listed in Supplemental Table S1.

### Luciferase assay

HEK293T cells were transfected with different pairs of pmirGLO reporter and miRNA-expressing plasmids or miRNA mimic. Dual-Luciferase^®^ reporter assays were carried out 48 hours post transfection according to the manufacturer’s instructions. Briefly, cells were lysed with 1X passive lysis buffer by gently shaking at room temperature for 15 minutes. Then, 20 µl lysate was added to 100 µl of Luciferase Assay Buffer II (LARII) and firefly luciferase activity was measured immediately. Finally, 100 µl Stop&Go reagent was added and Renilla luciferase activity was measured.

### Cell proliferation and migration assays

HEK293T, MDA-MB-231, and MCF-7 cells were seeded at a density of 0.3×10^5^ cells/well and transfected with synthetic miR-snaR or miR-snaR inhibitor or the appropriate control. To measure cell proliferation, the MTT (3-(4,5-dimethylthiazol-2-yl)-2,5-diphenyltetrazolium) assay was performed at 24 hours post transfection. The culture medium of each cell culture was replaced by 500 ng/µL MTT solution and incubated for 4 hours at 37°C. Following incubation, cell culture plates were centrifuged at 3,000 g for 10 minutes and supernatant was carefully removed. Formazan product was dissolved in 400 µl DMSO and the absorbance was measured across the UV spectrum from 570 nm to 690 nm with a plate reader. To measure cell migration, a confluent monolayer of cells was observed 24 hours post-transfection, at which time the monolayer was scraped with a rubber policeman and washed with 1X phosphate buffer saline to remove floating cells. Images were captured at 0, 24, and 48 hours after scraping. The wound healing size tool plugin for ImageJ was used to quantify wound healing at each timepoint.

## ACKNOWLEDGEMENTS

We thank Drs. Susan Frost, Shuo Gu, Narry V. Kim, Brian Law, and Jianrong Lu for sharing reagents; J. Bert Flanegan for critical reading of the manuscript; UF Interdisciplinary Center for Biotechnology Research and the UF Health Cancer Center Sequencing Core for sequencing services. This work was supported by grants from National Institutes of Health [R00-CA190886 and R35-GM128753 to M.X.; P01-CA214091 to R. R.; R01-CA188561 to R. O.], University of Florida Research Office [Research Opportunity Seed Fund to M.X.] and the University of Florida Informatics Institute [UFII Graduate Fellowship to D.S.]. Funding for open access charge: National Institutes of Health.

## AUTHOR CONTRIBUTION

D.S., R.R., and M.X. conceived the project. D.S. developed TOMiD and analyzed miR-snaR targeting. Y.L. characterized miR-snaR biogenesis. C.M.G. investigated miR-snaR targeting of NME1. L.L., C.J.F. and Y.L. predicted miR-snaR targets. C.J.F. analyzed small RNA-seq data. C.M.G. and P.N. performed NME1 western blots. P.N. and R.O. measured miR-snaR’s influcence on cell proliferation. D.S., Y.L., C.M.G., R.R. and M.X. wrote the manuscript.

## DATA AVAILABILITY

Ago-qCLASH data from Illumina high-throughput sequencing has been deposited in the Gene Expression Omnibus under accession number GSE164634.

