## supplemental methods for "A non-canonical microRNA derived from the snaR-A non-coding RNA targets a metastasis inhibitor"

**SUPPLEMENTAL MATERIALS AND METHODS**

**TOMiD analysis**

For each sample, 100-nt sequence reads were obtained utilizing the Illumina HiSeq3000 instrument at the University of Florida Interdisciplinary Center for Biotechnology Research. Sequences were transferred and stored on UF’s HiPerGator High-Performance Computing Environment and all subsequent analysis steps were performed on this system. Read quality control and analysis were first performed using FastQC (version 0.11.7) (Andrews and others 2010). Trimmomatic (version 0.36) was then utilized to remove adapter sequences and low-quality bases (PHRED <= 20) from read ends, with reads below 26 bases in length discarded from further analysis (Bolger et al. 2014). Reads from both mate pairs remaining after trimming were merged using PEAR (version 0.9.6) (Zhang et al. 2014) and the merged sequence files were combined with any unpaired reads resulting from trimming. Reads were sequenced utilizing a 4-nucleotide barcode to distinguish between PCR-duplicate and identical disparate reads. As such, PCR-duplicates were removed using fastx-collapser (fastx_toolkit, version 0.0.14) (Gordon and Hannon 2010), and barcodes were removed using cutadapt (version 1.8.1) (Martin 2011). Reads were then again collapsed using fastx-collapser, with count information tallying unique sequence reads encoded within read sequence identifiers (Gordon and Hannon 2010).

#FastQC Analysis:

$ fastqc -q --noextract --threads ${THREADS} ${READS_R1} ${READS_R2}

#Trimmomatic Trimming:

$ trimmomatic PE -threads ${THREADS} ${READS_R1} ${READS_R2} -baseout ${OUT_DIR}/{READS_NAME}.fastq ILLUMINACLIP:{WD}/reads_raw/RPI_trimmomatic.fasta:2:30:10 LEADING:3 TRAILING:3 SLIDINGWINDOW:4:15 MINLEN:26

#Mate Pair Merging via PEAR

$ pear -f ${READS_1P} -r ${READS_2P} -o ${OUT_NAME_MGD} -j ${THREADS} -y ${MEM_TOTAL}M -n 26

\# Collapsing of PCR-Duplicates via FASTX_Collapser:

$ fastx_collapser -v -i ${OUT_NAME_MGD} -o ${OUT_NAME_PCR_DEDUP}

\# Removal of Read Barcodes via cutadapt

$ cutadapt -u 4 -u -4 -m 18 -o ${OUT_NAME_PCR_DEDUP_CUT} ${OUT_NAME_PCR_DEDUP}

\# Collapsing of Quantitative Duplicates via FASTX_Collapser:

$ fastx_collapser -v -i ${OUT_NAME_PCR_DEDUP_CUT} -o ${OUT_NAME_PCR_DEDUP_CUT_QUANT_DEDUP}

For alignment of primary target sequences, the hOH7 reference sequence database included with the Hyb software package (version 20141126) (Travis et al. 2014) was utilized and an alignment reference index was created using Bowtie2-build (version 2.3.4.3) (Langmead and Salzberg 2012). Reads were aligned to the reference genome using Bowtie2 with parameters designed to allow local alignment of short segments of the reads to multiple reference sequences to identify possible read subsequences (Langmead and Salzberg 2012). Alignments were then sorted by query read name using Samtools sort (version 1.9) (Li et al. 2009).

#Alignment of Reads for Candidate Hybrid Identification
$ bowtie2 -f ${OUT_NAME_PCR_DEDUP_CUT_QUANT_DEDUP} -x ${INDEX_FILE} --threads ${THREADS} -S ${RAW_OUT_SAM} -D 20 -R 3 -N 1 -L 16 -k 20 --local -i S,1,0.50 --score-min L,18,0 --ma 1 --np 0 --mp 2,2 --rfg 5,1

#Sorting of Alignments:

$ samtools sort -n -o ${SORT_OUT_SAM} --threads ${THREADS} ${RAW_OUT_SAM}

Aligned reads for each sample were evaluated as potential hybrid sequences using a custom script implemented in Python3 using the pysam Python module with Samtools (Li et al. 2009; Heger et al. 2014; Van Rossum and Python development 2018). Alignments were first evaluated on a per-read basis ignoring all full-length alignments. For reads with partial-length alignments, the longer alignment for each read was selected as the most likely subsequence to represent the “target” of a chimeric RNA molecule. Reads with partial alignments were then identified as a candidate hybrid if the length of the unaligned portion of the sequence is between 18 and 23 bases and separated into a candidate miRNA (cmiRNA) and target sequence from the unaligned and aligned portions, respectively. The Gibbs Free Energy of hybrid formation between each pair of sequences was predicted using UNAFold hybrid-min (version 3.8)(Markham and Zuker 2008), with only candidate sequences with a predicted free energy value lower than -7.0 kcal/mol retained. Finally, each exact candidate miRNA sequence was required to have been called identically in two or more reads to reduce the effect of ligation and sequencing error on cmiRNA identification.

\# Analysis of Candidate Hybrids

$ CLASH_split_v10.py ${SORT_OUT_SAM} ‑o ${OUT_BASE} ‑‑script_mode ‑‑verbose ‑‑max_fold_energy "-7.0" ‑‑min_candidate_cov "2" ‑‑min_candidate_len "18" ‑‑max_candidate_len "23" ‑‑action_log ‑‑name_prefix ${READS_NAME}_cmiRNA_ > ${OUT_BASE}.out"

\# Prediction of Free Energy of Folding with UNAFold hybrid-min

$ hybrid-min -o ${OUT_BASE}.fold.UNAFold ${OUT_BASE}_5p.fasta ${OUT_BASE}_3p.fasta

cmiRNA molecules identified in sequences from each individual sample were then combined, duplicates were removed, and each candidate was assigned a unique identifier using a custom script implemented in Python3 (Van Rossum and Python development 2018). Each cmiRNA was required to exist in more than one sample for further processing, giving a total requirement of >= 4 total unique hybrids required for each cmiRNA to pass filtration. For identification of cmiRNA transcript source, the RefSeq human RNA reference library was acquired from ftp://ftp.ncbi.nlm.nih.gov/refseq/H_sapiens/annotation/GRCh38_latest/refseq_identifiers/GRCh38_latest_rna.fna.gz on 2019-03-19 and an alignment reference index was prepared via Bowtie2-build (version 2.3.4.3) (Langmead and Salzberg 2012; O'Leary et al. 2016). cmiRNA sequences were then aligned to the prepared reference transcriptome via Bowite2 (version 2.3.4.3) (Langmead and Salzberg 2012), and sorted via Samtools sort (version 1.9) (Li et al. 2009). A custom Python script utilizing pysam and Samtools was then used to annotate the candidate miRNA in each sample’s respective candidate hybrid sequences, and final hybrids were output in the Hyb file format (Li et al. 2009; Heger et al. 2014; Travis et al. 2014; Van Rossum and Python development 2018). Subsequent steps for visualization and quantification of identified cmiRNA and hybrid counts were then performed using additional scripts.

\# Combination of Sample-Specific cmiRNAs

$ CLASH_combine_v02.py ${IN_FASTAS} --out_name_prefix ${READS_NAME} -o ${OUT_BASE} --verbose --min_fasta_cov 2

\# Alignment of Combined cmiRNAs to RefSeq Database

$ bowtie2 -f ${CMB_FASTA} -x ${INDEX_FILE} --threads ${THREADS} -S ${OUT_SAM_RAW} -D 20 -R 3 -N 1 -L 16 --local -i S,1,0.50 --score-min L,18,0 --ma 1 --np 0 --mp 2,2 --rfg 5,1

\# Sorting of Alignments by Name

$ samtools sort -n -o ${OUT_SAM_SORT} --threads ${THREADS} ${OUT_SAM_RAW}"

\# Assignment of cmiRNA to Alignments

$ CLASH_assign_v03.py ${IN_HYB} -s ${OUT_SAM_SORT} -v --action_log -c ${CMB_CONTAINS} -o ${OUT_ASSIGN_BASE} --exclude_unclustered

**Identification of Drosha-independent miRNA**

For identification of possible Drosha-independent miRNAs, the counts of each detected cmiRNA within a unique hybrid were compared across the WT, Drosha-KO and Dicer-KO qCLASH datasets. The count of each cmiRNA per sample was first transformed into a proportion of that cmiRNA per sample by dividing the cmiRNA count by the total sample candidate hybrid count to normalize to the number of possible hybrids. A cutoff threshold was then applied retaining cmiRNA where $prop_{cmiRNA}\geq1.0*{10}^{-5}$ (approximately 25 reads/sample) to reduce the effect of large variations due to small sample counts, resulting in 672 remaining cmiRNA alignments. The qCLASH experiments providing sequencing data were conducted in two sets of two technical replicates. To reflect this distribution, category averages were first calculated across technical replicates per sample, and then again across different samples.

For each cmiRNA, Dicer-dependence was calculated as the proportion of the count of a specific miRNA in the Dicer-KO to its count in WT, and the Drosha-dependence was similarly calculated as the proportion in Drosha-KO compared to WT. A minimum Drosha-dependence score of ${dep}_{Drosha}\geq1.5$was set to require enrichment of a cmiRNA in the Drosha-KO condition. Although many cmiRNA had zero counts in the Dicer-KO condition, and thus a Dicer-Dependence score of ${dep}_{Dicer}=0.00$, a minimum Dicer-dependence value of ${dep}_{Dicer}=0.01$ was utilized in subsequent calculations to prevent errors from division by zero. A Drosha-independent miRNA discovery scoring metric was then calculated by dividing the Drosha-dependence by the Dicer-dependence, with a high score indicating Drosha-independence and Dicer-dependence for a given cmiRNA (Equations 1-3). A $score_{Drosha/Dicer}\geq20$ was chosen as a cutoff threshold to focus the analysis on the highest-scoring candidates (range 0 to 501).

$$prop_{cmiRNA}=\frac{count_{cmiRNA}}{count_{total}}$$

Equation 1: Calculation of cmiRNA Sample Proportion

$${dep}_{DRO}=\frac{prop_{DRO}}{prop_{WT}}, {dep}_{DI}=\frac{prop_{DI}}{prop_{WT}}$$

Equation 2: Calculation of cmiRNA Drosha-Dependence and Dicer-Dependence

$$score_{Drosha/Dicer}=\frac{{dep}_{DRO}}{{max(dep}_{DI},0.01)}, with {dep}_{DRO}\geq1.5$$

Equation 3: Calculation of combined Drosha/Dicer Site score from Drosha-Independence and Dicer-Dependence scores

**Identification of miR-snaR binding sites**

To identify the miR-snaR binding site(s) in all targets, miR-snaR reference ‘AGCCTGGTCCACATGGGTCGGA’ was added to hOH7 genome database to compose a customized database. Processed reads obtained from HCT116 WT and Drosha-KO cells were generated as mentioned above. Reads from 293 cells were download from GEO database (GSE46039) and processed as described (Helwak et al. 2013). We aligned processed reads with the customized hOH7 database by Hyb (version 20141126) and extended the sequences for every target by 25 nucleotides to ensure that the miRNA binding site is completely covered. Hyb analysis generated a viennad format file, which contains base-pairing pattern of the miRNA/target pair, the position and sequence of target in hOH7 database, and predicted folding energy. The position of the target was extracted from the viennad files. The miR-snaR target sites that include seed-region complementarity and appear in at least two biological replicates were identified for analysis in Fig. 4B. The start and end positions of the target are defined by the overlapping sequences in the reads mapped to the same site. We then composed a custom Python 2.7 script (Viennad_Call_Peak.py) to call miR-snaR peak region in three datasets (HCT116 WT, HCT116 Drosha-KO and 293 cells) based on the viennad files.

\# Align processed reads into hOH7 genome file and extend each target coordinateds by 25 nucleotides

$ hyb analyse fold=UNAfold in=${IN_FASTQ} db=${HOH7_GENOME} type=mim pref=mim

\# Identify miR-snaR binding site(s)

$ python2 Viennad_Call_Peak.py -i ${Viennad_file(s)} -g ${HOH7_GENOME} -r ${REPLICATE_FILE} -m {miRNA_name} -o ${output_name}

**miRNA/target interactions analysis**

To analyze the miR-snaR/target base-pairing pattern, the merged reads generated as described above were first filtered to contain only those containing miR-snaR utilizing Bowtie2 (Langmead and Salzberg 2012). Using the database that contains the annotated miR-snaR miRNA, Hyb was then utilized to identify CLASH hybrids from the filtered reads, including the use of UNAFold hybrid-min (version 3.8) for prediction of folding patterns (Markham and Zuker 2008; Travis et al. 2014). A Python3 script (hybkit_analyze_miR-snaR_folding.py) built on the hybkit API (version 0.2.1a) was utilized to filter reads containing mRNA or mRNA-pseudogene targets and calculate the per-base folding characteristics of miR-snaR (Stribling 2020).

\# Alignment/Selection of only miR-snaR-containing Reads:

$ LANG=C bowtie2 -f ${IN_FASTA} -x ${MIR_SNAR_INDEX_FILE} --threads ${THREADS} -S ${OUT_SAM} --al ${OUT_FASTA} --norc -a -R 3 -N 0 -L 16 --local -i S,1,0.50 --score-min L,18,0 --ma 1 --np 0 --mp 2,2 --rfg 5,1 --no-unal

\# Hyb miR-snaR Hybrid Identification

$ LANG=C BOWTIE2_PARAM=\"-D 20 -R 3 -N 1 -L 16 -a --local -i S,1,0.50 --score-min L,18,0 --ma 1 --np 0 --mp 2,2 --rdg 5,1 --rfg 5,1 -p ${THREADS}\" HYB_DB=${HYB_DB_DIR} hyb detect align=bowtie2 word=11 format=fasta analyse fold=UNAfold in=${IN_FILE} db=${DB_NAME}

\# Analyze miR-snaR Folding Pattern with Hybkit

$ python3 hybkit_analyze_miR-snaR_folding.py

To analyze the base pairing pattern of all miRNAs, merged reads were utilized directly with Hyb and UNAFold using the miR-snaR-augmented database(Travis et al. 2014). A Python3 script (hybkit_analyze_other_folding.py) built on the hybkit API (version 0.2.1a) was utilized to filter hybrids to only miRNA-containing hybrids without miR-snaR with targets of mRNAs or mRNA-pseudogenes. The per-base folding characteristics all non-miR-snaR miRNA in the dataset (Stribling 2020) was then calculated.

\# Hyb Hybrid Identification

$ LANG=C BOWTIE2_PARAM=\"-D 20 -k 200 -R 3 -N 1 -L 16 --local -i S,1,0.50 --score-min L,18,0 --ma 1 --np 0 --mp 2,2 --rdg 5,1 --rfg 5,1 --norc -p ${THREADS}\" HYB_DB=${HYB_DB_DIR} hyb detect align=bowtie2 word=11 format=fasta analyse fold=UNAfold in=${IN_FILE} db=${DB_NAME}"

\# Analyze non-miR-snaR Folding Pattern with Hybkit

$ python3 hybkit_analyze_other_folding.py

***In vitro* transcription of snaR-A**

To generate DNA templates, PCR reactions were performed in 50 µL with 1 unit of Phusion polymerase (NEB) with High Fidelity buffer, 40 nM forward and reverse primers, 200 nM dNTP, 10 U Phusion polymerase and 20 ng pBS-snaR-A. In a thermal cycler, repeat (98°C for 15s, 60°C for 20s, 72°C for 20s) for 35 cycles after initial denaturing 98°C for 30s. The DNA products were separated on a 3% agarose gel and then extracted using gel extraction kit (ZYMO research). Concentrations of PCR DNA were measured by Nanodrop. In a 20 µL in vitro transcription reaction, 250 ng of PCR DNA template, 20 U of Murine RNase inhibitor (NEB), 5 U of T7 RNA polymerase, 1mM (each) of NTP were incubated in 1X transcription buffer [40 mM Tris-HCl pH8.0, 25 mM NaCl, 2 mM spermidine(HCl)_3_ 8 mM MgCl_2_ and 10 mM DTT] at 37°C overnight and separated on a 6% denaturing PAGE gel. Extracted gels were soaked in RNA elution buffer (0.3 M NaOAc and 25 mM Tris) and incubated at room temperature overnight. RNAs were then PCA extracted from the elution buffer and measured concentration with Nanodrop.

***In vitro* Dicer cleavage assay**

Briefly, Flag-tagged human Dicers were overexpressed by transfection of the expression plasmid in HEK293T cells and purified by α-Flag antibodies. In cleavage assay, approximately 20 ng gel purified RNA was mixed with 2 µl 10 X buffer (200 mM Tris pH 6.5, 15 mM MgCl_2_, 250 mM NaCl, 10 mM DTT and 10% glycerol), approximately 100 ng Dicer on beads, and water was added to 20µl total volume. The mixture was incubated at 37°C for 1 hour with gentle agitation. The RNA was then PCA extracted and subjected to northern Blot analysis.

**Plasmid and miRNA mimic transfection**

Plasmid DNA was transfected using polyethylenimine (PEI). HEK293T cells were seeded 24 hours prior to transfection. PEI and plasmid DNA were separately diluted in OPTI-MEM (Gibco) and incubated at room temperature for 5 minutes. Diluted PEI and DNA were then combined and incubated at room temperature for an additional 15 minutes before being applied to cells. Cells were then incubated at 37°C. Cell medium was changed 24 hours post-transfection. Total RNA was collected 48 hours post-transfection. For transfection in 24-well plates, 1 µg of plasmid DNA and 4 µg of PEI were used per well. For 6-well plates, 2 µg of DNA and 8 µg of PEI were used per well.

miRNA mimic was transfected using Lipofectamine RNAiMAX transfection reagent. HEK293T, MCF7, and MDA-MB-231 cells were seeded immediately prior to transfection. RNAiMAX and miRNA mimic were separately diluted in OPTI-MEM and incubated at room temperature for 5 minutes. Diluted RNAiMAX and miRNA mimic were combined and incubated at room temperature for an additional 15 minutes before being applied to cells. Cells were returned to incubation at 37°C and harvested 48 hours post-transfection. For 6-well plates, 30 pmol of miRNA mimic and 9 µl of RNAiMAX were used per well. For 24-well plates, 5 pmol of miRNA mimic and 1.5 µl of RNAiMAX were used per well.

**Ago immunoprecipitation**

α-Ago antibody (clone 4F9) was used for Ago-IP from HEK293T, MCF7, MDA-MB-231 and HCT116 cells. Approximately 1 million cells were lysed by 200 µL of NP-40 lysis buffer (50 mM Tris–HCl pH 7.5, 1% NP-40, 10% glycerol, 150 mM NaCl, 5 mM EDTA, and 0.5 mM PMSF) for 30 minutes at 4°C and supernatant was collected by centrifugation at 21,000 × g for 10 minuets at 4°C. Total protein concentration in the supernatant was measured by DC protein assay kit (Bio-Rad). Total protein lysate was subjected to immunoprecipitation with α-Ago antibody-coated Protein L beads (Pierce, SE251848) for 2 hours at 4°C. After incubation, the beads were washed twice with cold Buffer D (20 mM HEPES pH 7.9, 10% glycerol, 0.1 M KCl, 1 mM EDTA, 0.1% NP-40 and 0.5 mM PMSF). RNAs were PCA extracted and subjected to Northern blotting analysis.

**Synthetic miRNA mimic**

Synthetic mature miRNA and miRNA* were obtained from Integrated DNA Technologies (IDT) and annealed to generate miRNA duplexes. The synthetic sequences were designed to contain a 5' phosphate on the mature miR-snaR strand (indicated by “/phosphate/”) and a 2-nucleotide 3' overhang. miR-snaR and miR-snaR* were combined with siRNA annealing buffer (100 mM potassium acetate, 2 mM magnesium acetate, 30 mM HEPES-KOH pH 7.4) and incubated for 1 minute at 90°C followed by 1 hour at 37°C. The final concentration of annealed miRNA was 20 µM. The negative control miRNA and miR-snaR antagomir (inhibitor) were designed and synthesized by GenePharma.

**Western blotting**

Total protein lysate was separated using 10 or 12 % sodium dodecyl sulfate polyacrylamide gel electrophoresis (SDS-PAGE). Contents of the SDS-PAGE were transferred to nitrocellulose or PVDF membrane using LifeTech transfer module at 0.3 Amp for one hour. The membrane was then incubated with 5% milk blocker in 1X TBST (150 mM NaCl, 20 mM Tris pH 7.5, 0.1% Tween-20) for one hour at room temperature. Next, the membrane was divided, and appropriate sections were incubated with antibody against NME1 (1:1000 diluted in 5% milk) or control proteins (GAPDH or Hsc70, 1:10,000 diluted in 5% milk) at 4°C overnight. Membrane sections were then washed 5 times with 1X TBST for 5 minutes each. Membrane sections were then incubated with HRP-conjugated secondary antibody IgG (1:5000 – 1:10,000) in 5% milk blocker in 1X TBST buffer for one hour at room temperature. This was followed by washing with 1X TBST five times for five minutes each. Finally, membrane sections were incubated with enhanced chemiluminescence (ECL) reagent mixture (1 part oxidizing reagent:1part luminol reagent) (Perkin Elmer) for 1 minute before being scanned on the Bio-Rad ChemiDoc imager.

**Northern blotting**

10 – 20 µg of total RNA from each sample was separated using 15% Urea-PAGE. Contents of the Urea-PAGE were transferred to Hybond N+ membrane (GE) using LifeTech transfer module at 0.2 Amp for one hour. The membrane was crosslinked twice using 254 nm UV crosslinker at 120 mJ/cm^2^. In the hybridization oven, membrane was incubated with 10 ml ExpressHyb hybridization solution in a hybridization tube for 30 minutes at 30°C. Membrane was then hybridized overnight at 30°C using probes conjugated with IR-dye (Table S1). Membrane was then washed twice. The first wash used 2X saline-sodium citrate (SSC)-0.1% SDS wash buffer. The second wash used 1X SSC-0.1% SDS buffer. For both washes, membrane was shaken at 110 rpm for 10 minutes at room temperature. Following washes, membrane was imaged on Amershan Typhoon scanner (GE health) to detect emission at 600 nm and 800 nm.

**SUPPLEMENTAL FIGURE LEGENDS**

**Supplemental Figure 1. Mapping the 5' and 3' ends of snaR-A transcripts.** Sequencing chromatogram of PCR products from 3' RACE (**A**) and 5' RACE (**B**) of snaR-A. (**C**) Schematic of sequencing results of individual clones obtained from 5' RACE of snaR-A.

**Supplemental Figure 2. SnaR-A and miR-snaR expression levels in HEK 293T Drosha-KO and HCT116 cells.** (**A**) Northern blot detection of snaR-A and U6 in total RNAs extracted from cells transfected with pBS or pBS-snaR-A. (**B**) Read number of miR-snaR in small RNA seq from HCT116 cells.

**Supplemental Figure 3. Multiple sequence alignment for the NME1 3' UTR in different primate species.** The sequence that is fully complementary with miR-snaR seed sequence is indicated by the bracket at the bottom. In the available representatives of the Homininae subfamily of higher-order primates: *Homo* (humans), *Gorilla* (gorillas), and *Pan* (chimpanzees and bonobos), absolute conservation of the complete miR-snaR binding site is observed (*Homo sapiens*, *Gorilla gorilla gorilla*, and *Pan paniscus)*. Absolute site conservation is also observed in *Pan troglodytes* (NM_001204512.1; not shown).

**Supplemental Figure 4. miR-snaR does not influence cell proliferation.** (**A**) Western blots show reduction of NME1 protein in 293T and MDA-MB-231 cells after transfection of miR-snaR mimic with GAPDH as a loading control. Asterisk (*) marks a non-specific band. (**B**) Left: Western blots show reduction of NME1 protein in MCF-7 cells with Hsc70 as a loading control. Asterisk (*) marks a non-specific band. Right: Quantification of three experiments as shown left. (Student’s t test, **p≤0.01). (**C**) Bar graph of MTT assay measurements (absorbance measured in the range of 570-690 nm) using cells transfected with either a control or a miR-snaR mimic. Error bars represent standard deviation from three experiments. Differences are not statistically significant (n.s., Student’s t test, p>0.05).

**Supplemental Table 1. Sequences of oligonucleotides used in this study.**
