## Supplementary material for "A non-canonical microRNA derived from the snaR-A non-coding RNA targets a metastasis inhibitor": fig. s1

### Figure S1

**A** Sanger sequencing of PCR products from 3′ RACE

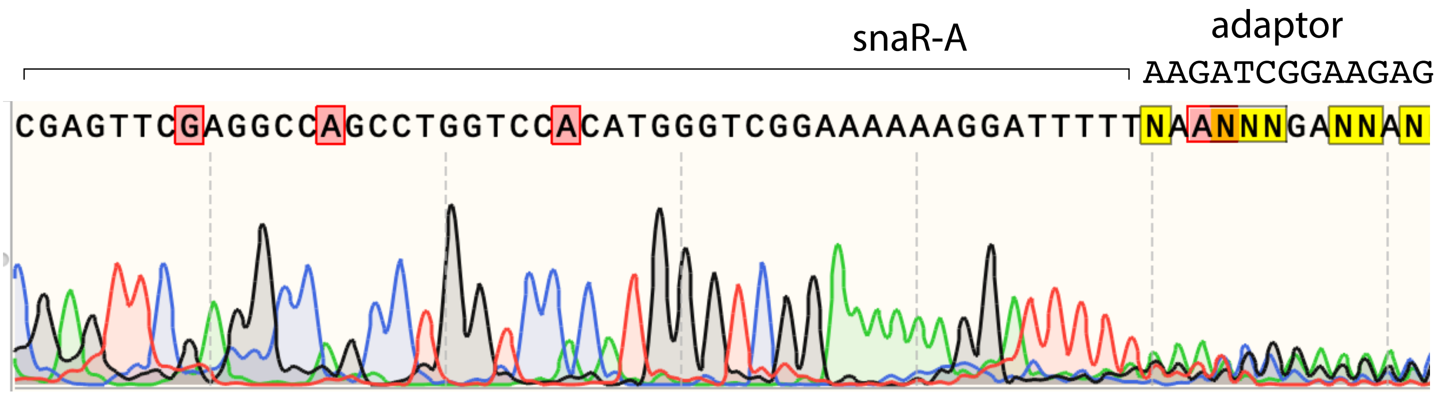


**B** Sanger sequencing of PCR products from 5′ RACE

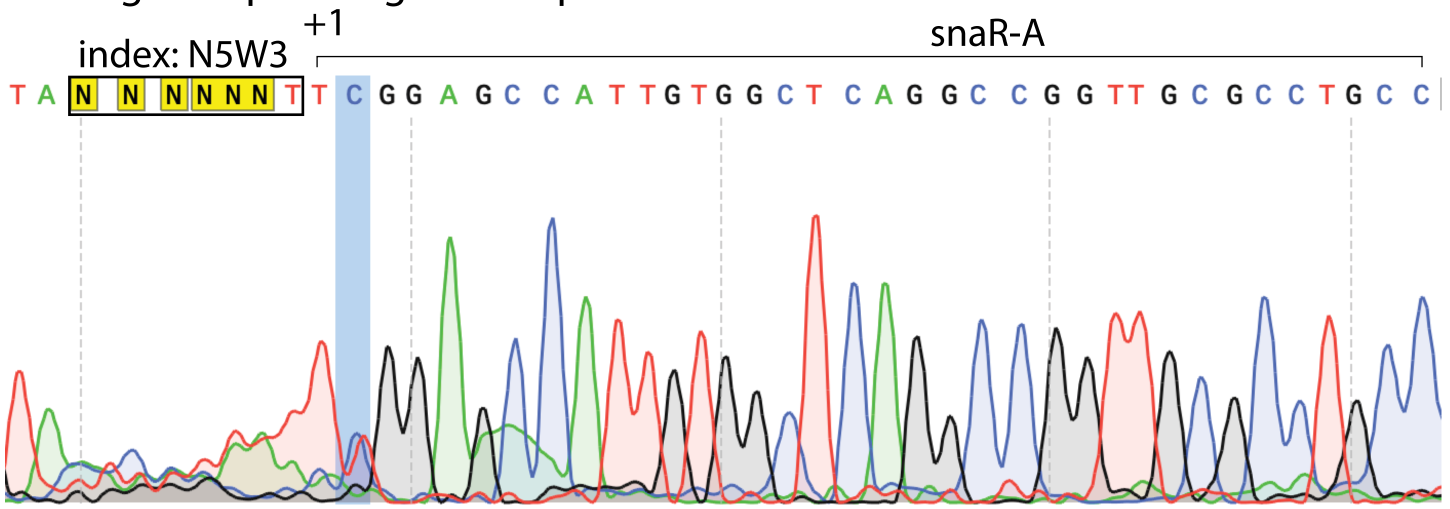


**C** Sanger sequencing of individual clones from 5′ RACE

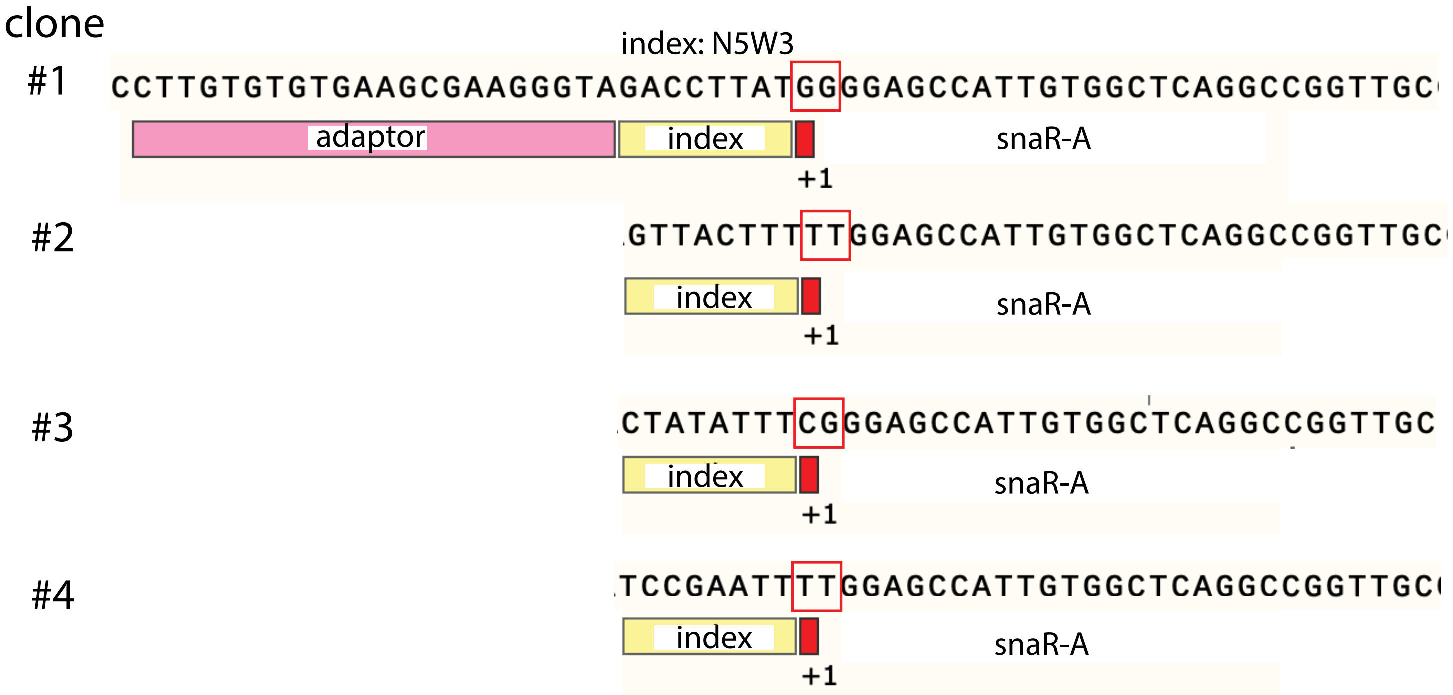
