## Supplementary figures and images for "A non-canonical microRNA derived from the snaR-A non-coding RNA targets a metastasis inhibitor"

### fig. s2

Figure S2

A

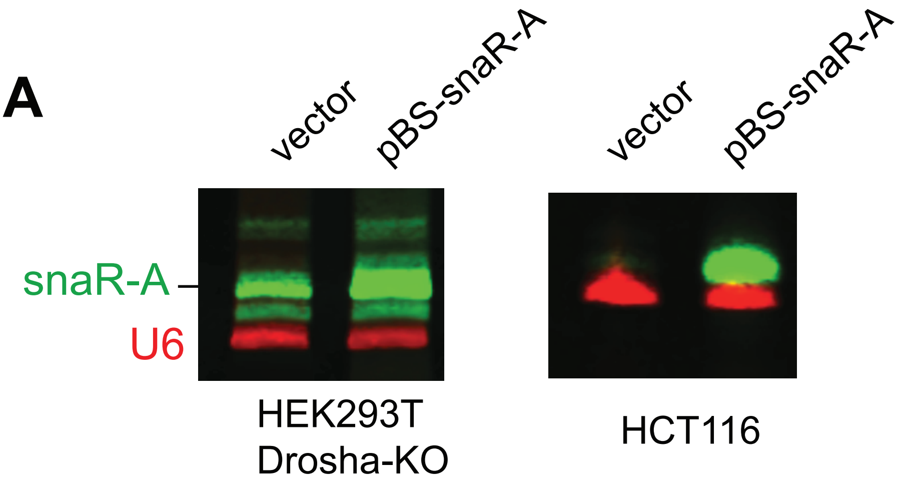


B

| HCT116 cells | miR-snaR read # |
|---|---|
| **WT** | 322 |
| **Drosha-KO** | 26,181 |
| **Dicer-KO** | 2 |
| **XPO5-KO** | 49 |

### fig. s3

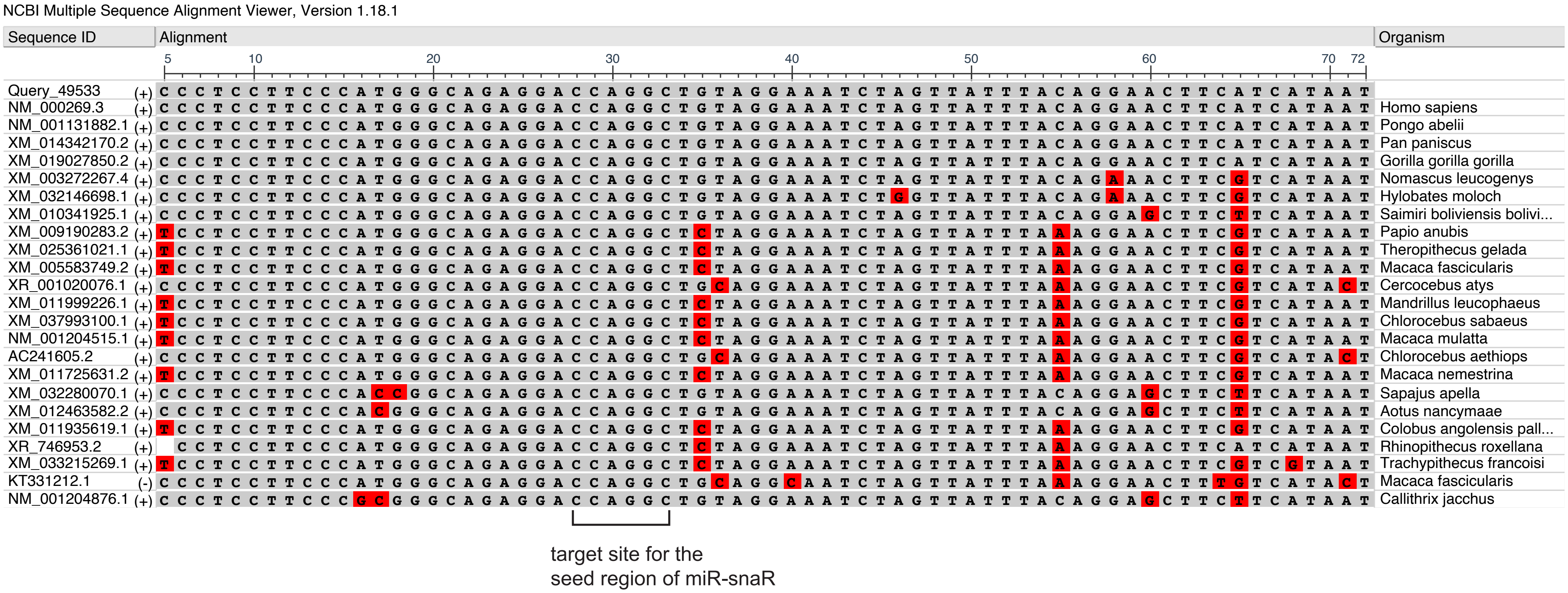

### fig. s4

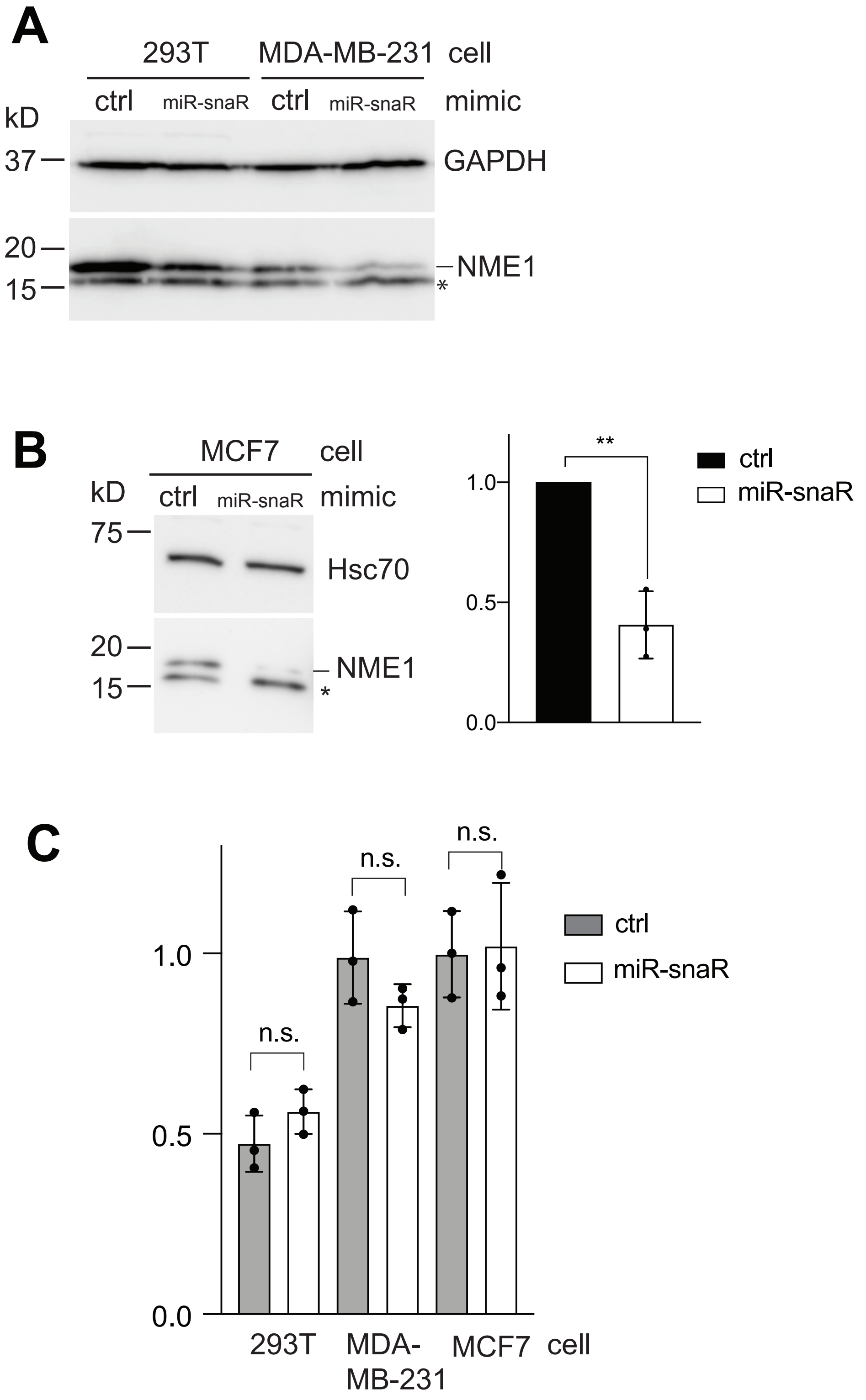
